# Synthesizing Mechanistic Hypotheses from Single-Cell Omics via Discretized Feature Attribution and Empirical Language Model Grounding

**DOI:** 10.64898/2026.07.09.737344

**Authors:** Jerry C.C. Chen, Yunqi Hong, Alexandra Bermudez, Jimmy K Hu, Cho-Jui Hsieh, Neil Y.C. Lin

## Abstract

Single-cell multimodal omics offer unprecedented resolution of cellular networks, yet translating continuous computational attributions into structured, testable biological mechanisms remains a persistent bottleneck. To address this limitation, we introduce an analytical pipeline employing decision trees to discretize continuous neural network attributions into explicit regulatory thresholds. These boundaries then structurally constrain large language models, enabling them to integrate established literature with empirical data to synthesize context-specific hypotheses. Applying this continuous-to-discrete framework across sparse datasets yielded novel biological mechanisms. Specifically, the framework articulated a cytoskeletal gating hierarchy governing EGF-stimulated pathways, identified transcriptomic drivers of input resistance in cortical interneurons, and delineated translational logic predicting Ki-67 abundance within spatial transcriptomics. Retrospective benchmarking validated the framework’s capacity to autonomously reconstruct published regulatory logic. Supported by a locally deployable open-weight language model and a code-free interface, this approach establishes an auditable methodology to extract robust experimental hypotheses from high-dimensional single-cell data.

## INTRODUCTION

Single-cell multimodal omics now enables the simultaneous measurement of epigenomic, transcriptomic, and proteomic states [1, 2]. While these multi-dimensional datasets provide the critical readouts to unbiasedly decode cellular decision-making [3, 4], translating them into actionable mechanistic hypotheses remains a key computational opportunity [5]. To navigate this complexity, linear transformations have established a robust standard for dimensionality reduction, successfully organizing omics data into interpretable patterns [6–11]. Building on these foundations, the field is increasingly shifting toward nonlinear models to capture the context-dependent dynamics that govern biological systems, such as threshold effects [12], saturation kinetics [13], and epistatic interactions [14].

Nonlinear models, including graph-based manifold learning [15] and ensemble regression [16] methods, have been successfully employed to reconstruct branching cell state trajectories. Furthermore, deep neural networks have been widely adopted to capture non-additive interactions within highly sparse datasets [17, 18], with self-supervised and transfer-learning extensions enabling the resolution of context-dependent biological structures and responses [19, 20]. As the field benefits from these unprecedented predictive capabilities, an emerging goal is to supplement this performance with mechanistic transparency [21]. While deep learning architectures implicitly capture complex biological features, translating their continuous mathematical parameters into explicit, rule-based regulatory mechanisms remains a research frontier [22, 23].

To further this goal, research has focused on developing attribution frameworks. Pioneer methods such as SHapley Additive exPlanations (SHAP) [24] and Integrated Gradients (IG) [25] attribute model predictions to input features (e.g., transcript levels of specific genes [21]). While these scores quantify continuous relative importance, they do not explicitly define the underlying combinatorial rules, such as nonlinear expression thresholds [26] or multi-gene gating [27]. Concurrently, large-scale biological foundation models, such as scGPT [28], GenePT [29], Geneformer [30], and Universal Cell Embedding [31], have shifted the computational paradigm toward generalized representations of cellular states. The encoding of these states into abstract vector spaces introduces an opportunity to develop extraction techniques capable of isolating the regulatory thresholds required to derive targeted wet-lab hypotheses [32, 33]. To facilitate this translation, researchers increasingly utilize large language models (LLMs) for literature synthesis [34, 35]. As LLMs increasingly serve as interpretability agents [36], recent studies stress the need to ground their outputs in data to prevent deceptive explanations [37, 38]. Simultaneously, integrations of multimodal data demonstrate how data-driven structural architectures can guide LLM-assisted biological annotation [39]. Building upon these seminal advances, the next opportunity is moving from broad functional annotation to targeted mechanistic understanding. This transition requires defining the thresholds where continuous predictive features become biological rules. Consequently, advancing interpretability depends on mapping continuous attributions to discrete, data-driven boundaries, thereby grounding LLM-generated hypotheses in verifiable mechanistic logic.

Here, we introduce Parsing Integrated-gradients and Trees to Create Hypotheses (PITCH), a computational framework that converts high-dimensional latent representations into explicit regulatory rules and literature-grounded hypotheses. PITCH models multimodal data using a Multilayer Perceptron (MLP) and applies IG to capture cell-specific feature attributions. A Decision Tree (DT) module then delineates cellular subpopulations, effectively translating these continuous importance scores into discrete decision thresholds [40]. To bridge computational outputs and biological interpretation, we structured a prompting framework to constrain an LLM for translating these DT-derived thresholds into mechanistic hypotheses. After validating the pipeline’s ability to recover underlying nonlinear logic on synthetic manifolds, we applied PITCH across three distinct biological paradigms. Specifically, the framework mapped structural parameters predictive of growth-factor signaling, identified transcriptomic signatures of electrophysiological variance in single neurons, and extracted transcriptomic boundaries governing spatial protein translation. To ensure accessibility, we deployed this pipeline as PITCH Studio, an interactive web interface that enables researchers to transition from retrospective observation to prospective experimental design.

## RESULTS

### A Continuous-to-Discrete Framework Recovers Simulated Regulatory Logic

Our post-hoc interpretability framework, PITCH, processes pre-normalized multimodal single-cell datasets, readily compatible with standard pipelines like Seurat or Scanpy for scRNA-seq, CITE-seq, and spatiogenomics, through a continuous-to-discrete analytical pipeline (Figure 1). The workflow begins by modeling cellular states, defined here as the multi-omic expression profiles (e.g., transcripts and surface proteins) of individual cells, using a two-layer MLP (blue box, Figure 1). This shallow neural network architecture was selected as an optimal intermediate. It provides sufficient nonlinearity to capture the epistatic, context-dependent molecular interactions, while deliberately avoiding the convoluted gradient landscapes of deeper networks that frequently destabilize post-hoc attribution.

**Figure 1.**
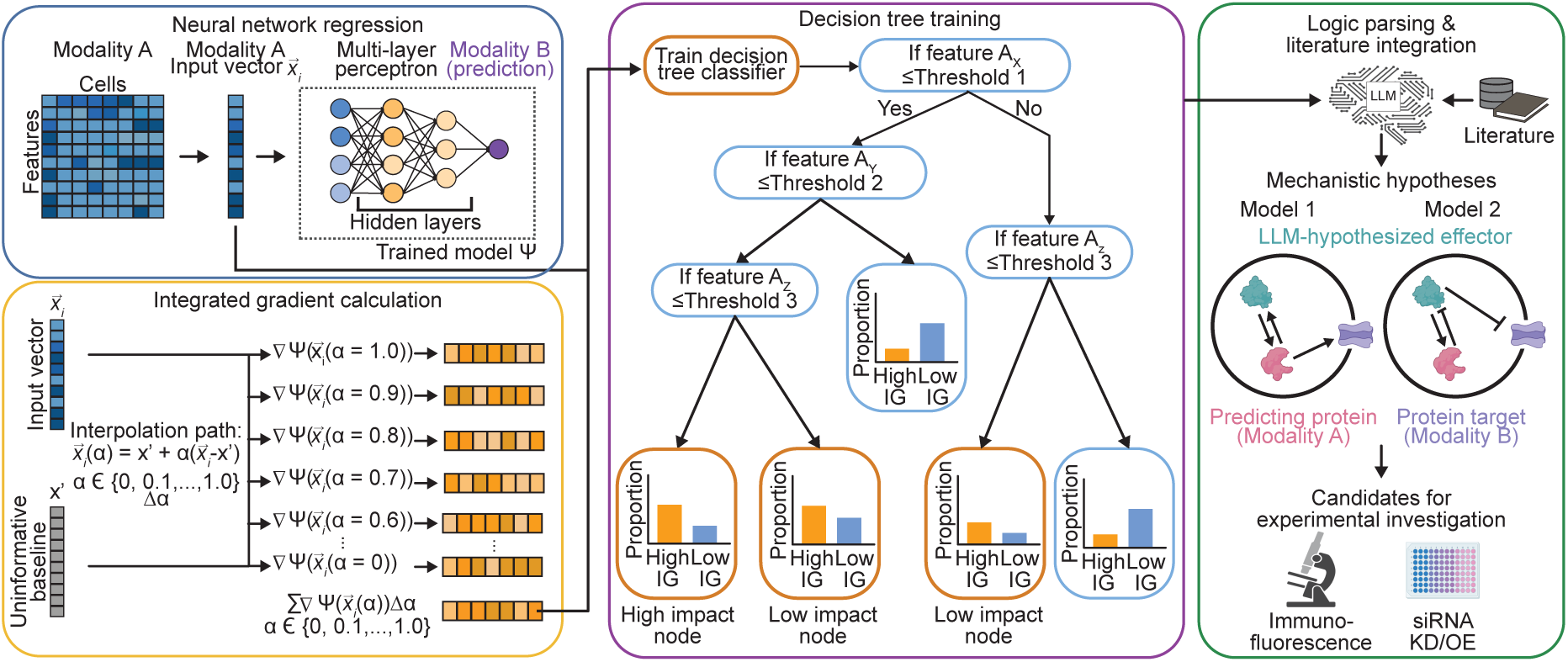
PITCH translates multimodal data into testable mechanistic hypotheses. PITCH bridges mathematical feature attribution and literature synthesis through a four-stage analytical pipeline. Stage 1 — neural network regression: Features from Modality A are encoded to predict a paired target response in Modality B using an MLP. Stage 2 (orange box) — IG calculation: To decode the MLP’s opaque latent space, IG extracts context-specific feature attributions by measuring gradients from an uninformative baseline to the observed single-cell state. Stage 3 (purple box) — decision tree training: A decision tree classifier translates these attribution scores into discrete regulatory boundaries, identifying the hierarchical feature conditions that define high-impact cellular subpopulations. Stage 4 (green box) — logic parsing and literature integration: An LLM synthesizes the extracted decision-tree logic with the literature to generate structured mechanistic hypotheses.

Within this latent space, biological features become mathematically entangled, obscuring their individual predictive roles. To decode these representations, we implemented a dual-stage extraction module. First, we use IG to calculate cell-specific attributions for each molecular variable (orange box, Figure 1). By mathematically tracing the model’s prediction backwards from a specific single-cell state to a neutral population baseline, IG provides a complete decomposition of how much each transcript or protein drove the final phenotypic prediction. Second, we train a DT classifier to threshold the highest attributions, isolating cellular subpopulations where the variable of interest exerts a dominant predictive effect (purple box, Figure 1). By recursively partitioning the multi-omic data space, the DT identifies the combinatorial molecular interactions most strongly associated with the targeted biological response. This non-parametric approach isolates the specific subpopulations correlated with a given biomarker, effectively translating abstract mathematical weights into hierarchical boundary conditions.

Recognizing that extracting mathematical rules is only half the challenge of biological discovery, we integrated an LLM to contextualize the DT-derived logic within established literature (green box, Figure 1). Serving as the final stage of the framework, this module integrates the extracted regulatory rules into explicit, testable mechanistic hypotheses for targeted wet-lab experiments.

We first validated the framework’s capacity to recover known predictive logic on a synthetic dataset with analytically defined ground truth. To simulate biological motifs such as additive baseline effects, non-monotonic concentration dependencies, and epistatic gating, we designed the function f = 2x_0_ – 3(x_1_)^2^ + 5*σ*(x_2_)*σ*(x_1_ + x_3_) (Figure S1A, top). After training the MLP to fit this manifold (R^2^ > 0.99, Figure S1A, bottom), IG attribution correctly identified x_1_ as the dominant negative driver, reflecting its amplified quadratic influence (Figure S1B). To delineate the conditions driving this attribution, we defined the most negative 10% of IG scores as the target class for DT classification. The resulting DT (n = 16, 000) partitioned the dataset, identifying that high-magnitude negative attribution is primarily driven by extreme values of x_1_ (x_1_ > 1.748 or x_1_ ≤ –1.607; Figure S1C, orange outline). Furthermore, the DT captured the underlying interaction terms, demonstrating that in non-extreme x_1_ regimes, the boundaries for strong attribution depend on x_3_ (e.g., –1.607 < x_1_ ≤ –1.595 and x_3_ > –0.431; Figure S1C, red outline). Finally, 2D input space plots confirmed that these DT-derived boundaries successfully encapsulate the regions of maximal negative attribution (Figure S1D).

### Nonlinear Modeling Predicts Context-Dependent pERK Signaling Dynamics

We next evaluated our pipeline’s ability to extract mechanistic regulatory logic from heterogeneous experimental biological observations. We tested a multiplexed signaling dataset [41] that records 12 dynamic signaling markers across a six-point titration of Epidermal Growth Factor (EGF; 0, 1, 6.25, 10, 25, and 100 ng/mL). To represent the preexisting cellular state, the data incorporates ∼ 500 curated parameters quantifying single-cell morphology, protein abundance, and local neighborhood context (Figure 2A). Because these structural and molecular profiles are unaffected by EGF stimulation, they serve as ideal baseline predictors of signaling dynamics. While previous analyses [41] established that EGF signaling outcomes are broadly predictable from these intrinsic states, the explicit combinatorial rules driving these predictions remain under-characterized (Figure 2A). Motivated by this gap, we aim to articulate the gating hierarchies governing EGF-induced signaling activity.

**Figure 2.**
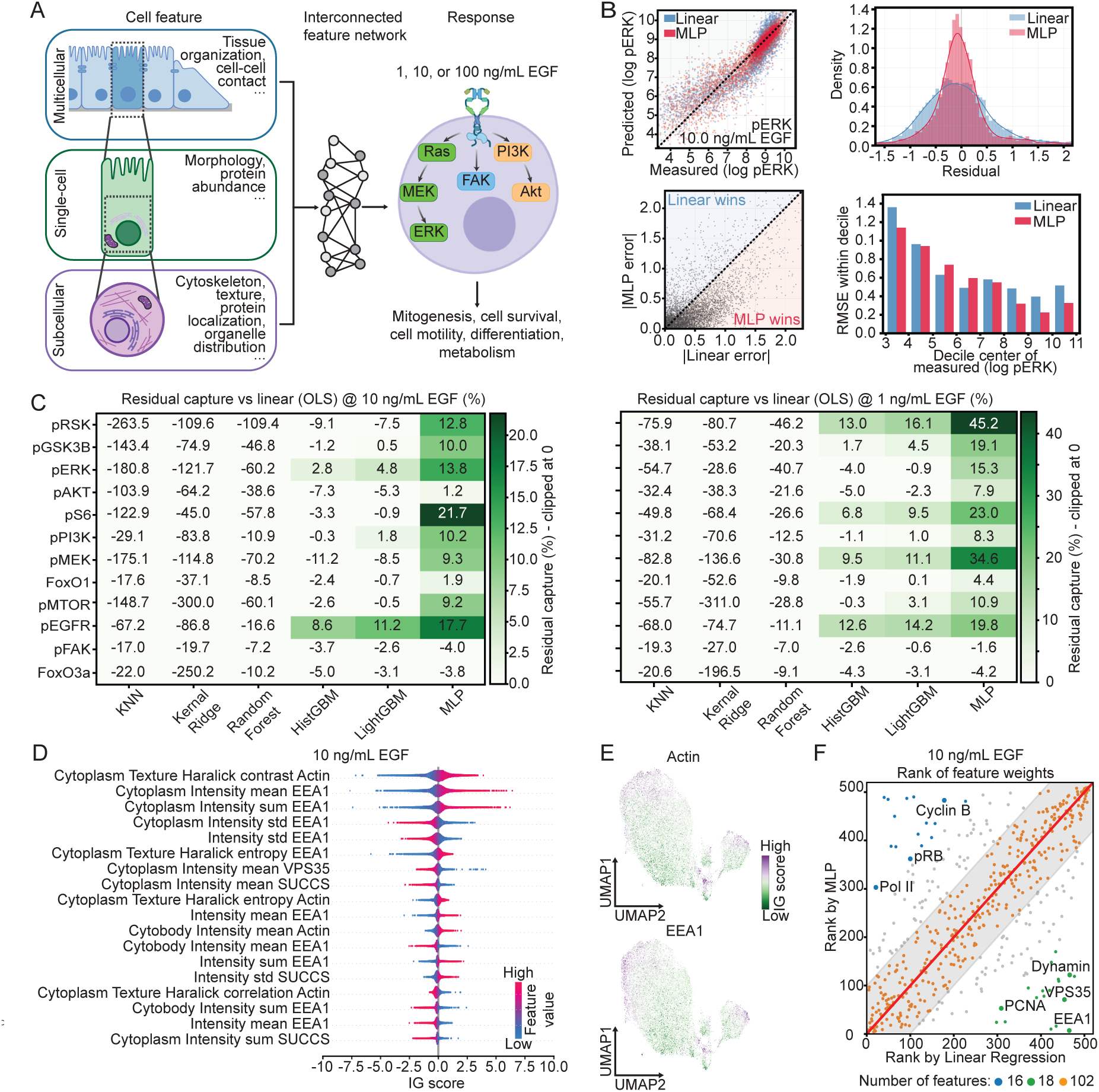
Coupling of MLP and IG reveals that actin and EEA1 are critical to pERK activity. (A) Multiscale cellular features (multicellular, single-cell, and subcellular) are quantified to define intrinsic cell states that govern downstream signaling and functional responses to EGF perturbation. (B) The MLP outperforms a baseline linear regression model in predicting pERK activity at 10 ng/mL EGF, demonstrating higher alignment with measured values, a narrower residual density centered at zero, and reduced absolute and root mean square errors (RMSE) across response deciles. (C) Benchmarking against five nonlinear regression models reveals the MLP achieves the highest residual capture (relative to an OLS baseline) across the majority of 12 signaling hubs at sub-saturating EGF concentrations (1 and 10 ng/mL). (D) Beeswarm plot of IG attributions for pERK prediction at 10 ng/mL EGF. Cytoplasmic Actin and EEA1 emerge as top predictive features, correlating positively with pERK activity. (E) UMAP projections colored by IG scores for features: intensity mean Actin (top) and intensity mean EEA1 (bottom). The attribution gradients highlight high context-dependency, while their concordant distributions suggest functional coupling. (F) Feature importance rank comparison between the MLP (ranked bottom-to-top) and linear regression (ranked left-to-right) at 10 ng/mL EGF. The MLP prioritizes receptor internalization machinery (e.g., EEA1, VPS35; green), whereas the linear model favors broad cell cycle markers (e.g., Cyclin B, Pol II; blue). Consistently ranked markers are shown in orange.

To capture dependencies across EGF concentrations, we implemented a unified MLP architecture modulated by a trainable encoder. By embedding the stimulus dose into a learned latent vector that scales internal representations, the framework leverages shared cellular features to isolate dose-specific regulatory dynamics. We observed that linear and nonlinear models performed comparably at saturating EGF concentrations (100 ng/mL) (Figure S2A). However, at sub-saturating levels (1 and 10 ng/mL), the MLP architecture proved superior. Linear regression of pERK levels at 10 ng/mL yielded systematic residuals, whereas the MLP better captured response variance (Figure 2B). We interpret this divergence as reflecting dose-dependent signaling dynamics. Consistent with established literature [42], saturating concentrations uniformly trigger kinase cascades, overriding preexisting cell state variations. Conversely, signaling at physiological thresholds is actively gated by internal states (e.g., organelle distribution or cytoskeletal tension) rather than guaranteed by receptor binding, necessitating the MLP’s nonlinear modeling capacity.

To systematically benchmark the MLP’s performance, we evaluated its residual reduction relative to a linear baseline across 12 response markers, comparing it against five representative nonlinear regression models (Figure 2C). The analysis confirmed that the MLP consistently achieved superior predictive accuracy for major signaling hubs, including pRSK, pERK, pS6, and pEGFR. These findings indicate that at sub-saturating concentrations characteristic of physiological stimulation, the coupling between intrinsic baseline features and primary signaling hubs is largely context-dependent. We focused our subsequent logic characterization on pERK, given its central role in phenotypic determination and its pronounced nonlinear dynamics within this regime.

### Predictive Molecular Attribution Identifies Context-Specific Correlates of pERK Activity

To unravel the regulatory rules associated with context-dependent signaling, we interrogated the trained MLP models using IG analysis to identify the key cellular determinants most predictive of single-cell pERK activity under 1 ng/mL and 10 ng/mL EGF stimulation conditions. Mean absolute IG attributions identified immunofluorescent intensities of actin (cytoskeleton) and EEA1 (endosomal trafficking) as prominent predictive components at both concentrations (Figures 2D and S2B). These findings implicate mechanotransduction and EGFR internalization as the primary biological axes governing ERK activity [43, 44]. Furthermore, the IG scores for both actin and EEA1 exhibited broad distributions, suggesting that their predictive importance for ERK activity is context-dependent across the cell population. Visualizing local attributions on a UMAP projection of the cellular phenotypic space revealed concordant distributions between actin and EEA1 (Figures 2E and S2C). This alignment points to a potential functional association between these components, which may relate to the actin cytoskeleton’s established role in scaffolding endosomal trafficking during ERK signaling.

To evaluate whether nonlinear modeling reveals biological associations missed by additive approaches, we compared MLP feature rankings against a linear model. For methodological consistency, linear feature importance was ranked by absolute coefficients to parallel the MLP’s mean absolute IG weights. We found that the linear model favored broadly distributed cell cycle markers (e.g., Pol II, pRB, Cyclin B) linked to population-wide baseline shifts. These additive features likely reflect general cellular states, such as proliferative capacity, that uniformly scale the overall signaling baseline. Conversely, the MLP highlighted receptor internalization machinery (e.g., EEA1, VPS35, Dynamin) (Figures 2F and S2D), which act as context-dependent bottlenecks governing the acute EGF response. Rather than implying one modeling approach is superior, this divergence merely highlights their complementary utility, in which linear models effectively map global phenotypic states, whereas nonlinear MLP attribution is required to isolate specific dependencies.

### Discretizing Attributions Extracts Mechanobiological Thresholds for pERK Prediction in Epithelial Cells

To translate IG attributions into discrete combinatorial rules, we trained a DT classifier on a curated set of 22 essential mechanobiology features (Table S1) to isolate subpopulations where actin strongly predicted pERK activity. To establish a reliable IG score threshold, we evaluated classification performance across the top 1% to 25% of cells. We optimized this threshold by balancing biological specificity (Gini impurity < 0.2) against robustness to single-cell dropout artifacts (Jaccard stability > 0.7). The top 10% to 15% bracket uniquely satisfied both criteria (Figure S3A), prompting us to select the 10% threshold for all subsequent analyses.

The classification conditions defined by the decision tree mapped the hierarchical molecular interactions that characterize high-attribution cellular states (Figures 3A, 3B, and S3B). Across both the 1 ng/mL and 10 ng/mL EGF conditions, parameters representing cell-substrate adherence (elevated Paxillin) and transcription activation (enriched Nuclear YAP) emerged as early decision nodes (i.e., top root splits) of the tree, indicating that high abundance of both markers drives the predictions of EGF-induced pERK activity. This identifies them as the primary markers used by the model to partition the high-attribution state, implicating that adherence and mechanosensory activation could be essential context for actin-mediated regulation. To verify identification robustness, we projected the DT-defined subpopulations onto the UMAP space, confirming their alignment with continuous high-actin attribution regions (Figures 3C and S3C). The extracted logic also captured dose-dependent morphological associations. At 1 ng/mL, the DT selected for symmetrical, non-polarized spreading, defined by thresholds for abundant focal adhesions and low elongation (Figure S3B). In contrast, the 10 ng/mL rules relied primarily on robust focal adhesions and YAP nuclear localization (Figure 3B).

**Figure 3.**
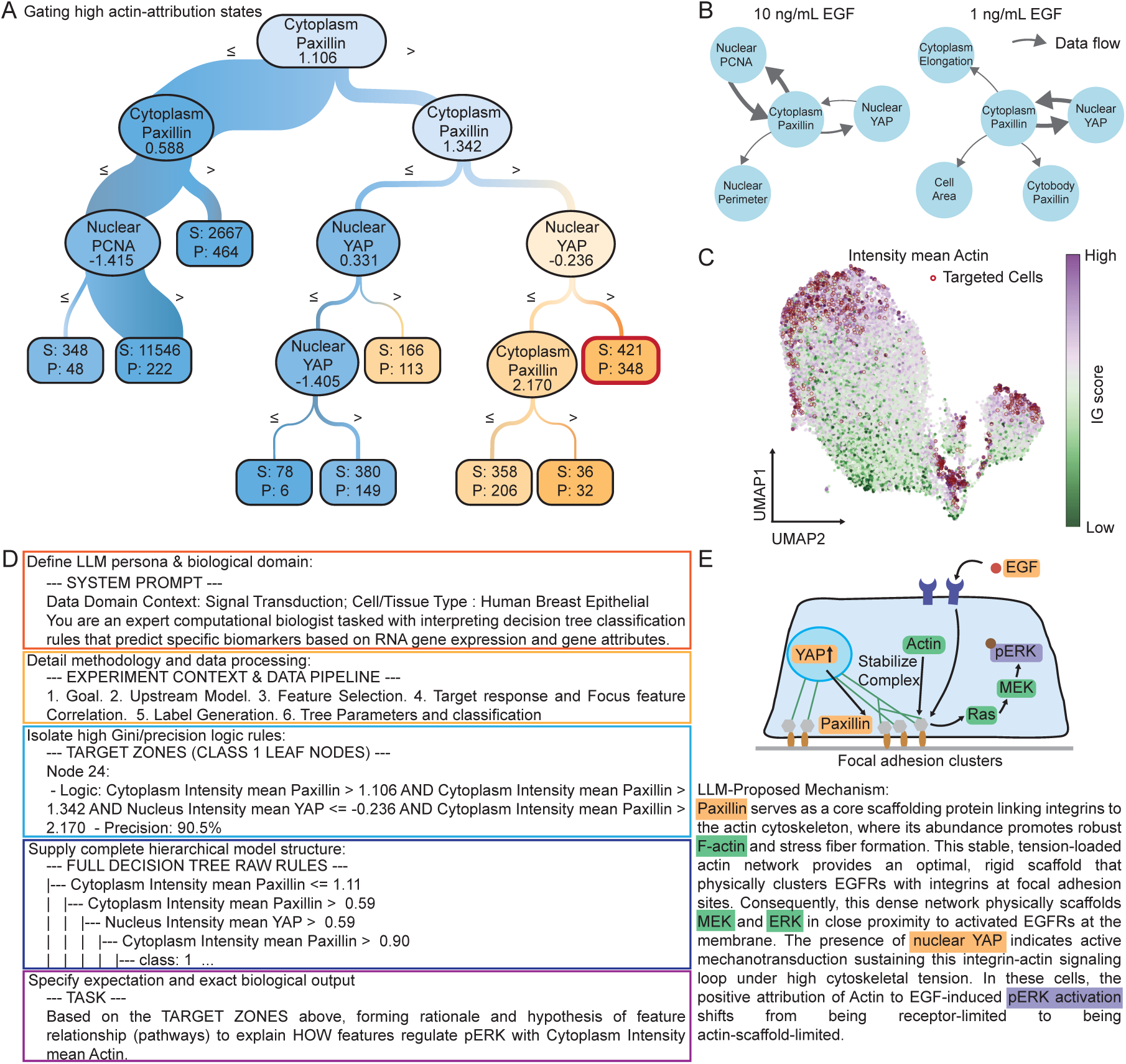
Integration of DT and LLM identifies mechanosensory feature hierarchies that regulate EGF-induced ERK phosphorylation. (A) A DT trained on 22 mechanobiological features maps the cellular hierarchy underlying high-actin attribution to pERK at 10 ng/mL EGF. Early splits on cytoplasmic Paxillin partition most of the population, while subsequent nuclear YAP splits isolate the highest-precision target subpopulation (red frame). Blue and orange nodes denote negative and positive classifications, respectively (S: total samples; P: positive samples). (B) Tree architectures reveal dose-dependent mechanobiological requirements: the high-actin attribution states rely primarily on mechanosensory biomarkers (e.g., Paxillin, YAP) at 10 ng/mL EGF, whereas broader morphological parameters at 1 ng/mL EGF (Figure S3B). Arrow width scales with sample size of the flow. (C) UMAP projection confirms that the DT-classified target subpopulation (red frame in (A)) aligns with continuous regions of high Actin attribution, corroborating the extracted decision rules. (D) Example of the structured LLM prompt used to translate DT logic into biological hypothesis. Integrating an expert persona, methodological context (orange), quantitative boundaries (yellow), statistical precision (light blue), DT results (deep blue), and explicit tasks (purple) grounds the LLM in empirical data to generate testable hypotheses. (E) Parsing the prompt inputs, the LLM formulates a hypothesis for the actin-pERK association: a tension-loaded actin network, anchored by Paxillin at focal adhesions, promotes the spatial colocalization of EGFRs with integrins and scaffolds the MEK-ERK cascade. Sustained by active mechanotransduction, this arrangement promotes pERK activation upon EGF perturbation.

To ensure these extracted rules were not methodological artifacts, we subjected the pipeline to two controls. First, the identified subpopulations remained consistent across three distinct IG baseline formulations (the global, 10 ng/mL local, and 0 ng/mL control averages) (Figure S4A). Second, applying the identical DT protocol to a negative control subset (the lowest 10% of actin IG scores) failed to converge, achieving only 53% precision (Figure S4B). This failure to model the negative control confirms that the extracted logic specifically captures actin’s predictive value rather than generic feature-selection bias.

### Generating Testable Hypotheses of Cytoskeletal Gating via Structurally Constrained LLMs

While DTs effectively isolate predictive subpopulations, manually synthesizing these discrete boundaries into cohesive, testable hypotheses is a time-consuming process that demands extensive domain expertise. We addressed this challenge by integrating the extracted logic into a structured LLM prompt (Figure 3D). By supplying the LLM (Gemini 3.1 Pro [45]) with exact quantitative boundaries, statistical precision metrics, and methodological metadata, we ground its reasoning in empirical data to synthesize biological rationales, design experiments, and provide literature references.

Applying this constrained framework to the 10 ng/mL EGF condition, the LLM proposed that within this epithelial context, mechanical tension promotes F-actin and stress fiber assembly, driving the spatial colocalization of EGFRs with integrins at paxillin-rich focal adhesions. While the LLM abstracts the precise molecular linkage, this proposed clustering aligns with established models [46] where the actin cytoskeleton orchestrates membrane compartmentalization, or shared lipid rafts, to trap these receptors together. This scaffolding facilitates the colocalization of downstream effectors (MEK and ERK), stabilizing the MAPK cascade and rendering EGF-induced signaling highly dependent on cytoskeletal integrity [47, 48]. To maintain this state of high cytoskeletal tension, the LLM implicates nuclear YAP in a mechanosensory feedback loop, where it sustains the paxillin-anchored focal adhesions required to stabilize the network. Consequently, the model translates the abstract DT criteria into a cell-type-specific prediction: targeted disruption of the actin scaffold will dismantle this shared biophysical hub, simultaneously suppressing YAP-driven mechanotransduction and acute receptor-driven pERK activation (Figure 3E and Figure S5). Notably, this autonomously generated hypothesis is corroborated by established empirical observations [44, 49–51].

To empirically test the model’s prediction regarding actin scaffold disruption, we examined transcriptional responses to pharmacological actin depolymerization using the LINCS L1000 database, which records high-throughput transcriptomic responses to diverse chemical perturbagens [52]. Consistent with the LLM’s prediction that actin integrity is physically required for upstream signaling, acute actin disruption (≤6 h) in receptor-driven epithelial cells simultaneously suppressed both MAPK activation and YAP/TAZ transcriptional targets (Figure S6, left). Crucially, in cell lines harboring active KRAS or PIK3CA mutations, which drive signaling downstream of the receptor, MAPK transcriptional output was unaffected by actin disruption, whereas YAP targets remained suppressed (Figure S6, right). This divergence corroborates the LLM’s mechanistic model: while downstream oncogenes can bypass the need for a membrane-proximal receptor scaffold, YAP activation remains critically dependent on actin scaffold integrity.

Finally, we conducted an ablation study to validate the prompt’s architectural design, confirming that these structural constraints are necessary for high-quality synthesis (Figure S7). Omitting experimental or statistical context severely degraded Biological Context Plausibility (a ∼5/10 drop) and Target-Feature Mechanism Fidelity (a ∼50% reduction), respectively. Furthermore, excluding extracted target zone candidates in favor of raw decision tree topologies crippled Zone Selection accuracy (∼2/10) and significantly impaired Cohesive Pathways Synthesis.

### Identifying Transcriptomic Predictors of Input Resistance in Cortical Interneurons

To evaluate PITCH’s generalizability across diverse biological domains, we applied the framework to three multimodal datasets encompassing patch-seq electrophysiology [53], image-based spatial transcriptomics [54], and CITE-seq surface profiling [55]. These datasets present distinct analytical challenges, including severe transcript dropout and cross-modal regulation. Across these varied contexts, PITCH demonstrated robustness to platform-specific artifacts (e.g., sparsity, noise).

We first applied PITCH to a single-neuron patch-seq dataset to predict the transcriptomic predictors of input resistance (R_in_). Capturing the nonlinear integration of these networks, the MLP accurately predicted R_in_ (Pearson’s r=0.77, Figure S8A). IG analysis recovered expected passive channels (*Kcnq5*, *Kcnmb4*, *Htr3a*) and identified *Fxyd6* (an active Na^+^/K^+^ ATPase regulator) as the top transcriptional predictor, proving the model captures multifactorial dependencies (Figure S8B). UMAP projection of *Fxyd6* IG scores then isolated its predictive importance to specific Caudal Ganglionic Eminence (CGE) (*Htr3a*^+^/*Cnr1*^+^/Cholecystokinin [*Cck*]^+^) and Medial Ganglionic Eminence-Somatostatin (MGE-SST) (*Lhx6*^+^/SST^+^/*Coro6*^+^) interneurons (Figure 4A). The DT partitioned these gradients into logic, defining an *Htr3a*^+^/*Cnr1*^+^ CGE-CCK subclass and a *Kcnq5*^−^/*Pde1a*^+^/*Coro6*^+^ MGE-SST subclass via combinatorial thresholds (Figure 4B, left; Figure S8C).

**Figure 4.**
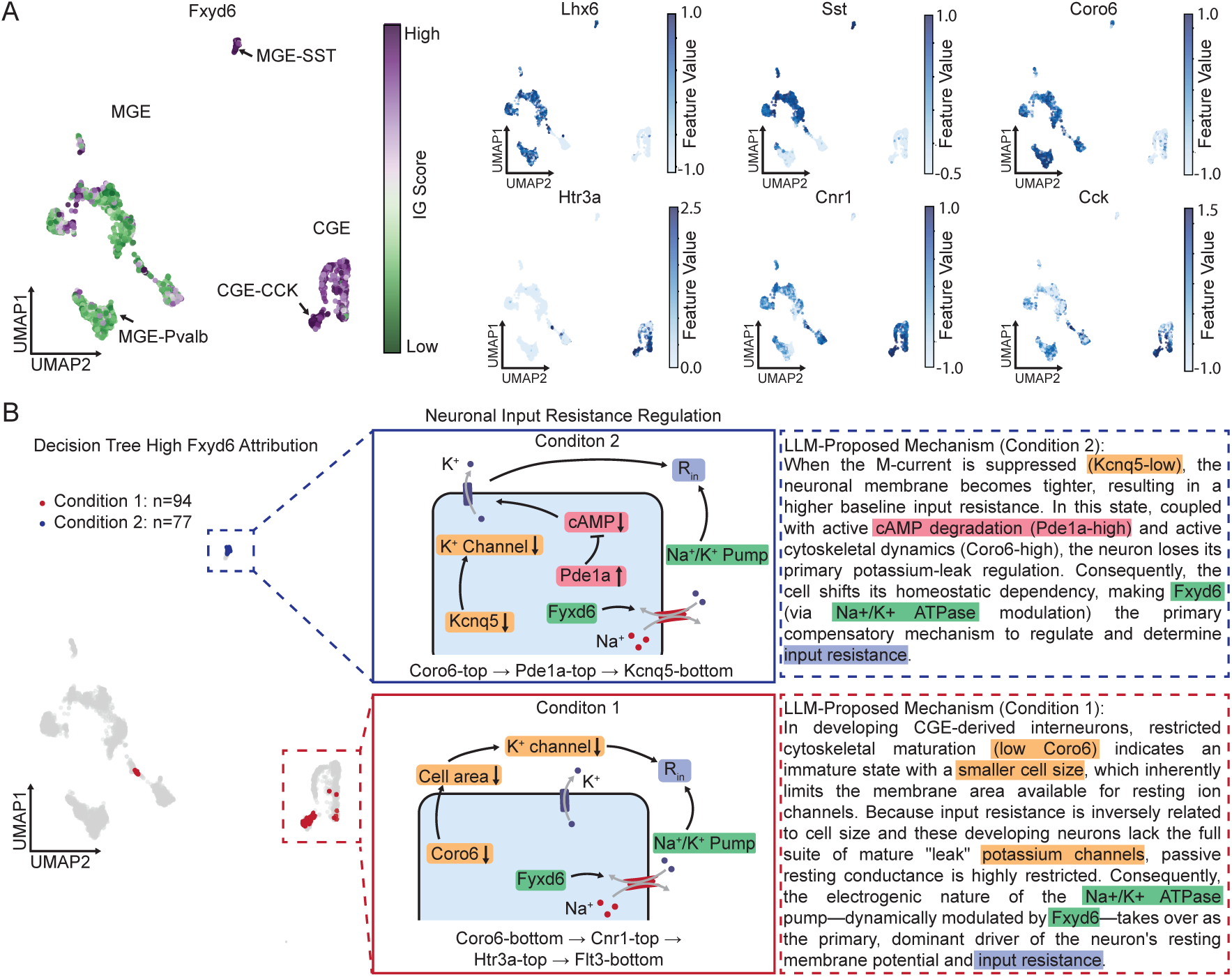
PITCH proposes state-dependent mechanisms of input resistance regulation in interneuron subpopulations. (A) UMAP projection of *Fxyd6* IG scores reveals high attribution in MGE-SST (*Lhx6*, *Sst*, *Coro6*) and CGE-CCK (*Htr3a*, *Cnr1*, *Cck*) interneuron phenotypes. (B) The LLM synthesizes state-specific combinatorial rules to propose that the *Fxyd6*-R_in_ association is driven by the suppression of primary ion leak currents. In the CGE-CCK lineage (Condition 1, red), downregulated actin-remodeling (*Coro6*) restricts cell size and passive potassium conductance, creating a high-R_in_ state reliant on *Fxyd6*-modulated Na^+^/K^+^ pumping. Conversely, in MGE-SST interneurons (Condition 2, blue), suppressed M-currents (*Kcnq5*) and elevated cAMP degradation (*Pde1a*) limit K^+^ leak channels, similarly shifting R_in_ dependency to the Na^+^/K^+^ pump. (Suffixes “-top” and “-bottom” denote expression above or below DT thresholds).

The LLM then synthesized these rules into de novo hypotheses (Figure 4B, right; Figure S9). For the CGE-CCK subclass (dependent on low *Coro6*), the LLM proposed that restricted cytoskeletal maturation, resulting in a smaller cell size, and fewer passive K^+^ channels maintain high baseline R_in_ [53, 56], maximizing sensitivity to *Fxyd6*-mediated active pumping [57]. For the SST state, defined by altered cyclic adenosine monophosphate (cAMP) and K^+^ channel profiles (*Pde1a*^+^/*Kcnq5*^−^) [58, 59], the LLM interpreted this “sealed leak” profile [60] as shifting R_in_ dependence toward active transport for electrophysiological homeostasis. While these overarching hypotheses represent a novel synthesis, each underlying mechanistic component is independently supported by the referenced literature.

### Extracting Hierarchical Predictive Logic from Sparse Spatial Transcriptomics

To assess robustness against data sparsity and post-transcriptional gating, we evaluated the pipeline’s ability to predict single-cell Ki-67 protein levels from breast cancer spatial transcriptomics [54]. Ki-67 is a canonical marker of active cellular proliferation, and its quantified protein abundance is clinically critical for grading tumor aggressiveness and guiding treatment decisions. By pairing multiplexed RNA detection with localized protein imaging, this dataset provides an ideal testbed for cross-modal prediction. Specifically, univariate profiling is frequently confounded by *MKI67* transcript dropout, and multivariable linear models struggle to capture the post-transcriptional gating that dictates final protein execution (Pearson’s r=0.44 and high residual error, Figure S10A, red) [61–63]. Overcoming these limitations, the MLP estimates single-cell Ki-67 abundance with higher accuracy (Pearson’s r=0.59 and residuals ∼ 0, Figure S10A, green). When subjected to random feature masking to emulate extreme data sparsity, the MLP consistently outperformed the linear baseline; even with 70% feature loss, it maintained a Pearson’s r of 0.44, matching the linear model’s peak accuracy on intact data (Figure S10A). IG attribution revealed that rather than relying on sparse *MKI67* transcripts, the network prioritized the broader translational machinery, identifying the chromatin-remodeling factor *PTMA* as the main predictor of Ki-67 (Figure S10B).

Projecting *PTMA* attribution onto the transcriptomic UMAP space mapped its predictive importance to a phenotypic gradient transitioning toward a proliferative mesenchymal-like state (*CKS2*^+^/*PGRMC1*^+^; Figure S10C). Using the IG scores, the DT delineated a distinct mitotic subpopulation defined by the co-expression rule: *RPLP0*^+^/ *HNRNPA2B1*^+^/ *TUBB4B*^+^ (Figure S10D). Structurally constrained by these data-driven boundaries, the LLM hypothesized a hierarchical regulatory axis (Figure 5A and Figure S11): upstream epigenetic remodeling (*PTMA*-driven chromatin decondensation [64]) links to coupled mRNA stabilization (*HNRNPA2B1*-mediated transcript binding [65]) and downstream translational execution (driven by *RPLP0*, a core 60S ribosomal protein), all operating within a mitotically active (*TUBB4B*^+^) state. Consequently, the LLM proposed this coordinated multi-omic cascade as the putative functional mechanism facilitating the execution of Ki-67 protein translation.

**Figure 5.**
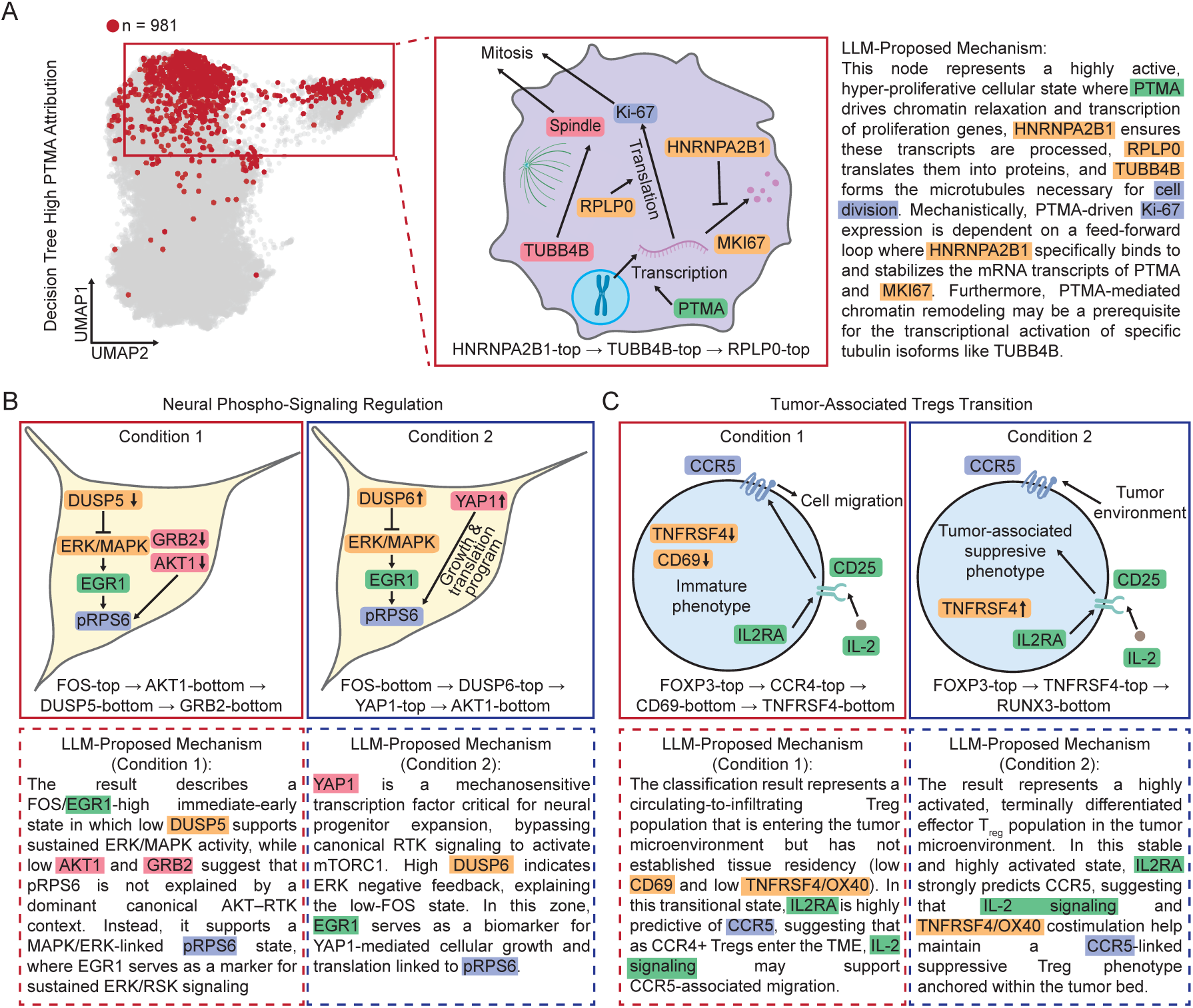
PITCH proposes epigenetic-to-translational hierarchies in breast cancer and retrospectively infers mechanisms across unseen datasets. (A) UMAP highlights a target subpopulation (n = 981) defined by high *PTMA* attribution. Synthesizing discrete decision tree boundaries, the LLM maps an integrated epigenetic-to-translational cascade linking *PTMA* (epigenetic remodeling), *HNRNPA2B1* (transcript binding), and *RPLP0* (translational execution) to collectively drive *TUBB4B*-associated mitosis and Ki-67 protein translation. (B) PITCH extracts context-dependent regulatory relationships for the translational hub pRPS6 (top 20% *EGR1* IG threshold). In a high-*FOS* context (Condition 1), the model delineates an alternative MAPK/ERK pathway where constrained *DUSP5* sustains *EGR1* activity independent of *AKT1* signaling. Conversely, a low-*FOS* context (Condition 2) reveals a compensatory bypass where an elevated *YAP1*/*TEAD* partnership drives local translation to maintain pRPS6 despite *DUSP6*-mediated feedback. (C) Modeling *CCR5* surface abundance in regulatory T cells (top 20% *IL2RA* IG threshold) reconstructs recruitment and maturation dynamics. Condition 1 defines a transitional, tumor-homing state (elevated *FOXP3*/*CCR4*, constrained *CD69*/*TNFRSF4*) where *IL2RA*-associated signaling supports *CCR5*-linked migration. Condition 2 isolates a terminally activated, tumor-resident state (elevated *FOXP3*/*TNFRSF4*, constrained *RUNX3*) where inflammatory *IL2RA* and *TNFRSF4* stimulation maintain a *CCR5*-linked suppressive phenotype. (Suffixes “-top” and “-bottom” denote expression above or below DT thresholds).

To evaluate the functional importance of these nominated targets in sustaining a proliferative phenotype, we analyzed genome-wide CRISPR knockout screens from the Cancer Dependency Map (DepMap) across 51 breast cancer models [66]. Single-gene essentiality scores demonstrated their varying dependencies. Specifically, the translational effector *RPLP0* proved essential for viability (mean Chronos = –1.52), whereas the chromatin-remodeling factor *PTMA* exhibited lower essentiality (mean Chronos = –0.45) (Figure S12A). Furthermore, bulk transcriptomics across these models indicated a coordinated regulatory module, revealing that *HNRNPA2B1* co-varies with both the mitotic marker *TUBB4B* (r = +0.79) and the translation effector *RPLP0* (r = +0.55) (Figure S12B). Collectively, these results confirm that PITCH effectively extracts co-regulated modules critical for cellular proliferation from single-cell snapshots. Beyond spatial transcriptomics, we further demonstrated the framework’s cross-modal versatility by predicting surface CD45RA abundance from paired CITE-seq transcriptomes [55], a task that requires the model to identify indirect regulatory proxies, as single-cell transcriptomics cannot easily resolve CD45 alternative splicing. The MLP captured CD45RA variance (Pearson’s *r* = 0.80, Figure S13A), with IG prioritizing cytotoxic machinery, specifically, cytolytic effectors (*GNLY*, *GZMB*) and their regulatory surface receptors (*KLRD1*), over canonical naive markers (Figure S13B). While standard UMAP failed to resolve these phenotypes (Figure S13C), the DT identified mature cytotoxic (*KLRD1*^+^/*NKG7*^+^/*GZMB*^+^/*CST3*^−^) and primed pre-effector (*NKG7*^+^/*GZMB*^−^/*S100A12*^−^) states (Figure S13D, S13E). Integrating this logic, the LLM hypothesized a developmental trajectory from early priming to terminal effector differentiation. We validated this predicted divergence by projecting the DT rules onto an independent peripheral blood mononuclear cell (PBMC) cohort (*n* = 2, 638). The derived boundaries partitioned cytotoxic NK and CD8^+^ cells from resting CD8^+^ T cells (Figure S14A, S14B) and confirmed the expected decoupling of *GNLY* and *GZMB* accumulation across the primed and mature states (Figure S14C).

### Retrospective Reconstruction of Published Regulatory Logic

As a definitive validation of our pipeline, we assessed its capacity to recover published biological mechanisms using unseen data modalities across two recent publications. Crucially, both studies were published after the training data cutoff of the utilized LLM (Gemini 3.1 Pro [45]), ensuring that the model could not rely on memorized knowledge. First, we analyzed a Phospho-seq dataset [67] designed to connect state-specific signaling activation to gene regulatory networks. The original study focused on phosphorylated Ribosomal Protein S6 (pRPS6) to map how converging kinases (including mTOR and ERK) coordinate with transcription factors to regulate translation. To evaluate if our framework could reconstruct this signaling logic, we predicted *RPS6* protein abundance solely from transcriptomic (scRNA-seq) features. By coupling MLP models with IG, we extracted gene-specific attributions (e.g., *EGR1*) (Figure S15A). DT partitioning and LLM-based evaluation successfully recovered the study’s core findings (Figures 5B, S15B, S15C and Table S2): an alternative *ERK*/*MAPK*-driven *RPS6* signaling axis (characterized by high *FOS*, with constrained *AKT1* and *DUSP5*) and a compensatory *YAP1*/*TEAD* regulatory partnership (high *YAP1* and *DUSP6*, low *FOS*).

Second, we modeled the recruitment and maturation dynamics of regulatory T cells (Tregs) in a hepatocellular carcinoma microenvironment [68]. Relying strictly on paired transcriptomic and surface protein profiles, we isolated Tregs using a *FOXP3*-independent marker panel to prevent target leakage and modeled *CCR5* surface abundance. IG identified *IL2RA* as a focal predictor of *CCR5*-associated migration (Figure S16A). The subsequent decision tree partitioning and autonomous evaluation reconstructed the previously reported progression from static data (Figures 5C, S16B, S16C and Table S3): a transitional, tumor-homing state (elevated *FOXP3* and *CCR4*; constrained *TNFRSF4* and *CD69*) supported by *IL2RA*-associated signaling, and a terminally activated, tumor-resident state (elevated *FOXP3* and *TNFRSF4*; constrained *RUNX3*) maintained by inflammatory signaling.

### PITCH Studio: A Code-Free Interface for Accessible Logic Extraction

While PITCH effectively translates single-cell data into mechanistic hypotheses, its reliance on advanced programming expertise limits widespread adoption. To address this, we developed PITCH Studio (https://linlabucla-pitch-studio.hf.space), a browser-based, code-free user interface that structures the workflow from initial model training to biological logic extraction (Figure 6A). Users upload preprocessed single-target, multi-omic data. Following MLP regression and molecular attribution, a selection module enables users to prioritize markers for downstream analysis. This interactive function is essential because biological networks are hierarchical, and uncovering secondary predictive layers requires iteratively shifting the analytical focus. By default, PITCH Studio assigns the highest-attribution marker as the primary target and selects the top thirty remaining variables as inputs, reducing dimensionality to isolate core subnetworks. Nevertheless, users can override these data-driven defaults to test specific biological hypotheses.

**Figure 6.**
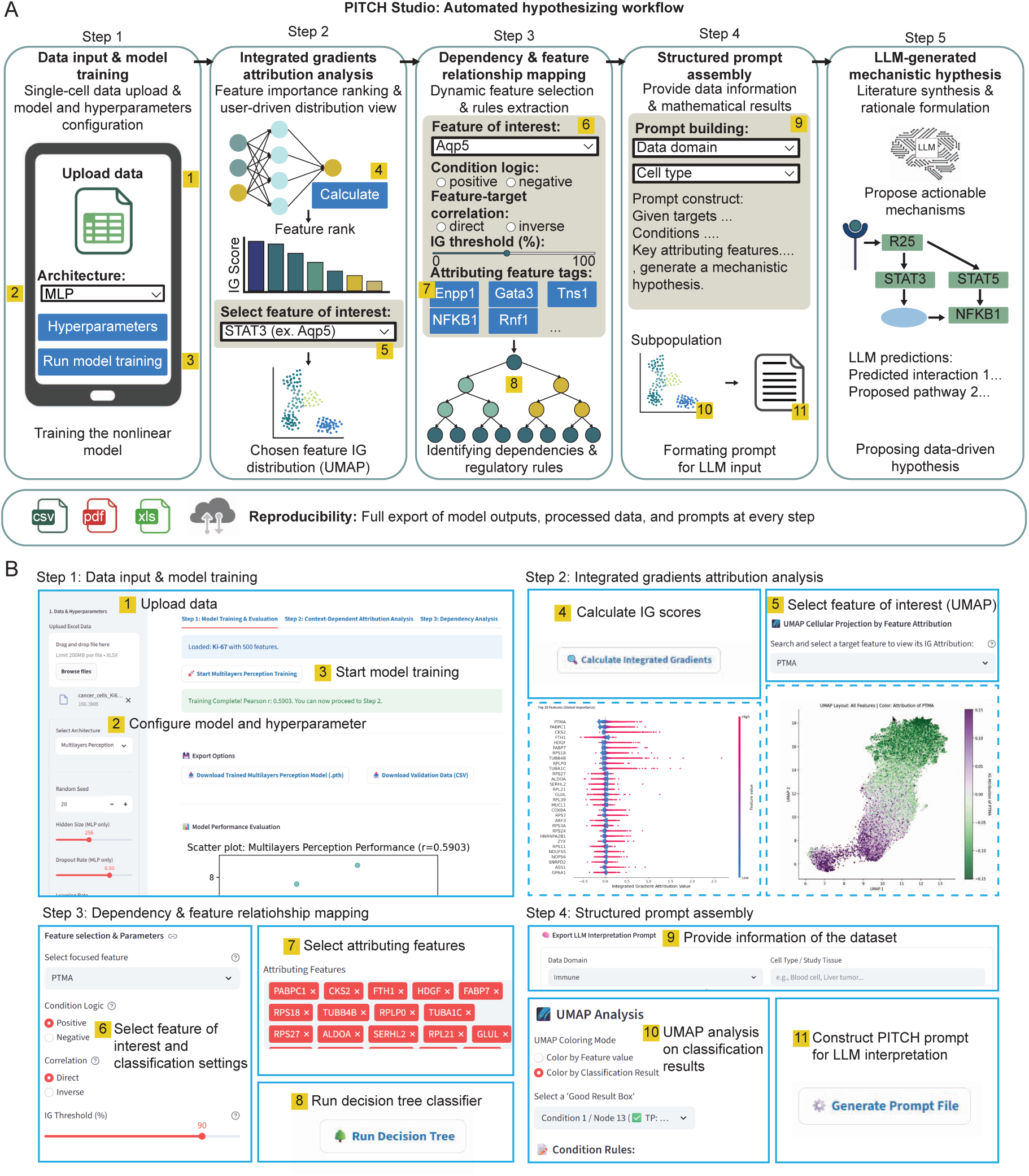
PITCH Studio: a code-free interface for automated multimodal analysis and mechanistic hypothesis generation. (A) PITCH Studio architecture and automated hypothesizing workflow. The browser-based platform guides users through a step-by-step analytical process, seamlessly integrating cross-modality model training (Step 1), feature attribution extraction (Step 2), and decision tree classification (Step 3). This pipeline culminates in a structured prompt builder (Step 4) that allows users to automatically formulate biological mechanisms via LLMs (Step 5). (B) User journey and operational guidance. The workflow begins by (1) uploading a dataset, (2) configuring hyperparameters, and (3) executing model training. The platform then (4) calculates Integrated Gradients, enabling users to (5) visualize target feature attributions via UMAP. To probe specific dependencies, users (6) define the target directional relationship and (7) select contextual features to (8) train a decision tree classifier. In the final interpretive phase, users (9) provide biological context and (10) evaluate classifications via orthogonal UMAP projections, allowing for (11) the automated generation of structured text prompts for downstream LLM interpretation.

To aid interpretation, the user interface integrates automated visualization and prompt-generation tools (Figure 6B). Users can visualize high-attribution subpopulations via UMAP and assemble structured LLM prompts to evaluate the underlying biological logic. Standardizing the prompt architecture minimizes user variance, ensuring reproducible translation of mathematical importance into mechanistic rationales. Finally, to maintain transparency and computational reproducibility, users can export all intermediate data and model outputs at any step of the workflow.

### A Locally Deployable Language Model Ensures Reproducible Hypothesis Synthesis

The final stage of PITCH utilizes an LLM to translate DTs into semantic reasoning. Relying on proprietary Application Programming Interfaces (APIs) introduces vulnerabilities, as model updates can induce prompt drift, yielding inconsistent biological interpretations over time. To ensure computational reproducibility and eliminate subscription barriers, we performed supervised fine-tuning (SFT) on an open-weight LLM, creating a locally deployable model to insulate future hypothesis generation from commercial API drift. While this student model may inherit the baseline literature biases of its teacher, it provides a version-controlled interpretive tool that ensures long-term reproducibility.

To generate training data, we simulated synthetic decision trees by systematically sampling cell types, markers, and pathway topologies to mimic the combinatorial complexity of authentic experimental outputs (Figures 7A and S17A). We prompted a high-capacity teacher model (Gemini 3.1 Pro [45]) to generate both final mechanistic hypotheses and internal reasoning steps. Incorporating these chain-of-thought trajectories encourages the model to sequentially bridge statistical thresholds with molecular functions before drawing conclusions. This dual-component dataset was used to SFT a 14B parameter student model (Ministral 3 [69]), scaled for execution on consumer-grade GPUs.

**Figure 7.**
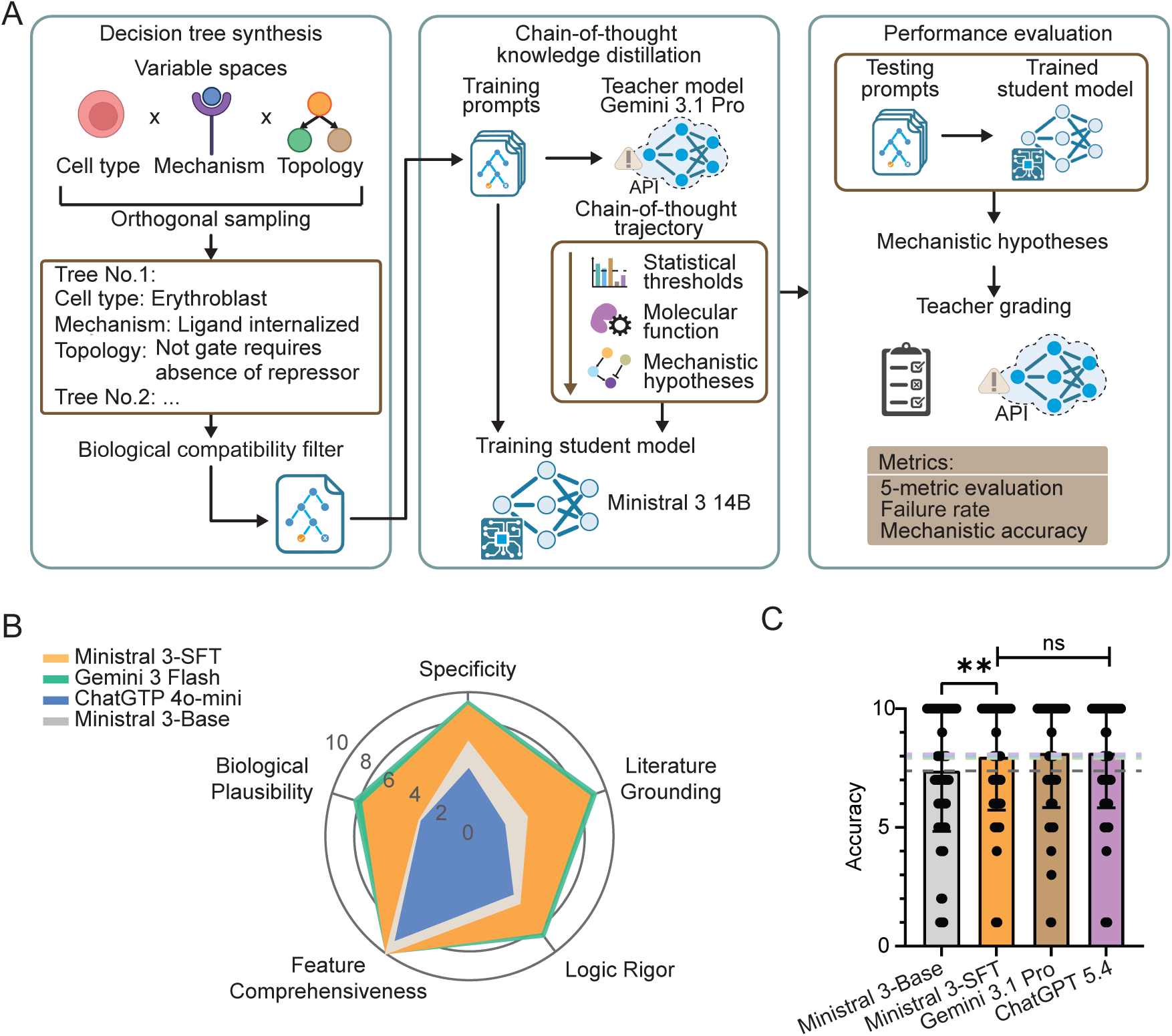
Training a local tree-interpretation large language model provides reliable reproducibility. (A) Schematic of the tree-interpretation model training pipeline. Synthetic decision trees are generated via orthogonal sampling across cell type, mechanism, and topology domains, followed by biological compatibility filtering. During chain-of-thought knowledge distillation, a teacher model (Gemini 3.1 Pro) processes these trees to generate reasoning trajectories and final hypotheses. These outputs then supervise the training of a local student model (Ministral 3 14B), which is subsequently evaluated by the teacher on unseen synthetic data. (B) Radar chart benchmarking model performance on a 10-point scale across five evaluative metrics: Specificity, Literature Grounding, Logic Rigor, Feature Comprehensiveness, and Biological Plausibility. While all evaluated models successfully retain contextual variables (high Feature Comprehensiveness), ChatGPT 4o-mini and Ministral 3-Base underperform in the remaining metrics (2–6 points). In contrast, the fine-tuned Ministral 3-SFT matches the performance of Gemini 3 Flash, consistently scoring 8–10 points across all domains. (C) Accuracy evaluation demonstrates that the fine-tuned Ministral 3-SFT significantly outperforms its base model, achieving mechanistic reconstruction accuracy comparable to advanced commercial models (Gemini 3.1 Pro and ChatGPT 5.4).

We evaluated the student model on a held-out synthetic dataset using a 10-point rubric assessing literature grounding, logical rigor, biological plausibility, and mechanistic specificity. At optimal checkpoints, the fully trained Ministral 3-SFT (average 8.4/10) demonstrated highly competitive performance against commercial models (Gemini 3 Flash, 8.9/10) across all criteria (Figure 7B and S17B). Crucially, optimal performance requires high structural precision.

When supplied with quantitative boundaries from the DT module, the SFT model accurately resolved ground-truth logic, matching the teacher model’s accuracy despite a ∼100-fold reduction in parameters (Figure 7C). By providing a high-performance local model, PITCH ensures data-driven hypothesis generation remains insulated from commercial API drift, providing an auditable interpretive framework.

## DISCUSSION

Balancing predictive capacity and mechanistic interpretability remains a central objective in modeling single-cell multi-omics [22, 70]. While methodologies that embed interpretability directly, such as Neural Additive Models [71], symbolic regression [72–74], or sparse mixture-of-experts [75, 76], are highly effective, they often necessitate trade-offs between capturing higher-order epistasis and maintaining computational scalability. To navigate these trade-offs, PITCH utilizes post-hoc, model-level rule extraction, adapting classical methodologies (e.g., TREPAN [77]) for high-dimensional omics. By projecting the learned latent space into decision boundaries via axiomatic gradient attribution, PITCH achieves mechanistic transparency without compromising predictive expressiveness.

Beyond extracting discrete predictive rules, PITCH integrates this logic directly into hypothesis generation. While LLMs and biological foundation models provide immense utility through generalized knowledge synthesis and broad cellular representations [30, 34, 78, 79], matching this global knowledge to the state-specific variance of an individual dataset remains critical for discovery [80]. By embedding empirically derived boundaries into the prompt, PITCH confines the language model’s reasoning to the mathematical gradients of the trained network. This approach complements generalized foundation models by supplying localized, dataset-specific predictive context. By translating multi-omic profiles into defined experimental directives, this framework narrows the search space for mapping therapeutic vulnerabilities. Such precision is poised to optimize the identification of epistatic drug synergies in multiplexed chemical transcriptomic screens [81], and it synergizes directly with latent-space perturbation modeling [82] by offering a mechanistic rationale for state-specific therapeutic resistance.

This capacity to generate highly localized directives aligns with a central objective of systems biology: identifying previously uncharacterized, context-specific dependencies within established networks. PITCH navigates this spectrum between canonical literature and novel observation. This capacity to discover context-specific biology is demonstrated by our retrospective analysis of the Phospho-seq dataset. Operating blinded to the original study’s conclusions, PITCH integrated established kinase biology with data-derived feature thresholds to deduce the dual-predictive relationships that constituted the paper’s core discoveries. By interpolating between existing knowledge and empirical variance, our framework provides a robust approach to formulating testable extensions of current biological understanding.

However, our framework operates within several constraints. First, PITCH inherently extracts predictive, rather than causal, feature attributions. While our retrospective benchmarking against independent CRISPR and pharmacological screens supports the functional and causal relevance of the extracted rules, deploying PITCH to map entirely uncharacterized pathways will ultimately require prospective experimental validation to confirm direct molecular mechanisms. Second, the reliance on IG [25] requires a continuous input space. This requirement makes the attribution process sensitive to the extreme sparsity and zero-inflation common in single-cell transcriptomics [5, 83]. Third, converting continuous attributions into discrete thresholds creates a structural trade-off. Limiting the depth of the decision tree preserves the clarity needed for the language model to parse the rules, but it restricts the pipeline’s ability to model higher-order interactions involving four or more molecular targets [70]. Finally, the mechanistic synthesis is fundamentally bounded by the language model’s training corpora [80]. This reliance introduces a systematic literature bias toward well-documented signaling pathways, rendering the framework less effective at proposing mechanisms for poorly characterized targets. Furthermore, as a pure text-reasoning engine, the LLM struggles to reliably resolve physical gene-to-complex mappings. Future iterations of the pipeline can address this by integrating structured external reference databases like CORUM [84].

Looking forward, a central utility of single-cell multi-omics lies in the construction of predictive *in silico* cell simulators, or cellular digital twins [85–87]. As these models are increasingly tasked with simulating complex pathologies and therapeutic responses, their underlying architectures require auditable predictive scaffolding rather than opaque associations. By serving as a modality-agnostic interface that extracts actionable rules from latent spaces, PITCH establishes a foundational step toward this goal. Ultimately, embedding explicit mechanistic transparency into computational frameworks will be essential for deploying neural networks across translational oncology and automated discovery pipelines.

## DECLARATIONS

### Author Contributions

Conceptualization: J.C.C.C., Y.H., C.J.H., and N.Y.C.L.; data curation: J.C.C.C. and Y.H.; investigation: J.C.C.C., Y.H., A.B., J.K.H., C.J.H., and N.Y.C.L.; methodology: J.C.C.C., Y.H., C.J.H., and N.Y.C.L.; writing – original draft: J.C.C.C., Y.H., A.B., and N.Y.C.L.; writing – review and editing: J.C.C.C., Y.H., A.B., J.K.H., C.J.H., and N.Y.C.L.;

## Acknowledgements

We thank Bernhard A. Kramer for guidance in understanding the data structure of the EGF experiment and for valuable comments on our regression results. We also thank Alexander Hoffman and Eric Deeds for insightful discussions and feedback on the pipeline. This work was partially supported by the National Science Foundation (CBET-2244760, DBI-2325121, IIS-2048280) and the NIH National Institute of General Medical Sciences (R35GM146735).

## STAR ⋆ METHODS

### KEY RESOURCES TABLE

#### Lead Contact

Neil Y.C. Lin

Cho-Jui Hsieh

#### Materials Availability

This study did not generate new unique reagents.

#### Data and Code Availability

- This paper analyzes existing, publicly available data. The accession numbers and source links for the datasets are listed in the key resources table.
- All original code has been deposited at GitHub and is publicly available as of the date of publication. The link is listed in the key resources table.
- Any additional information required to reanalyze the data reported in this paper is available from the lead contact upon request.

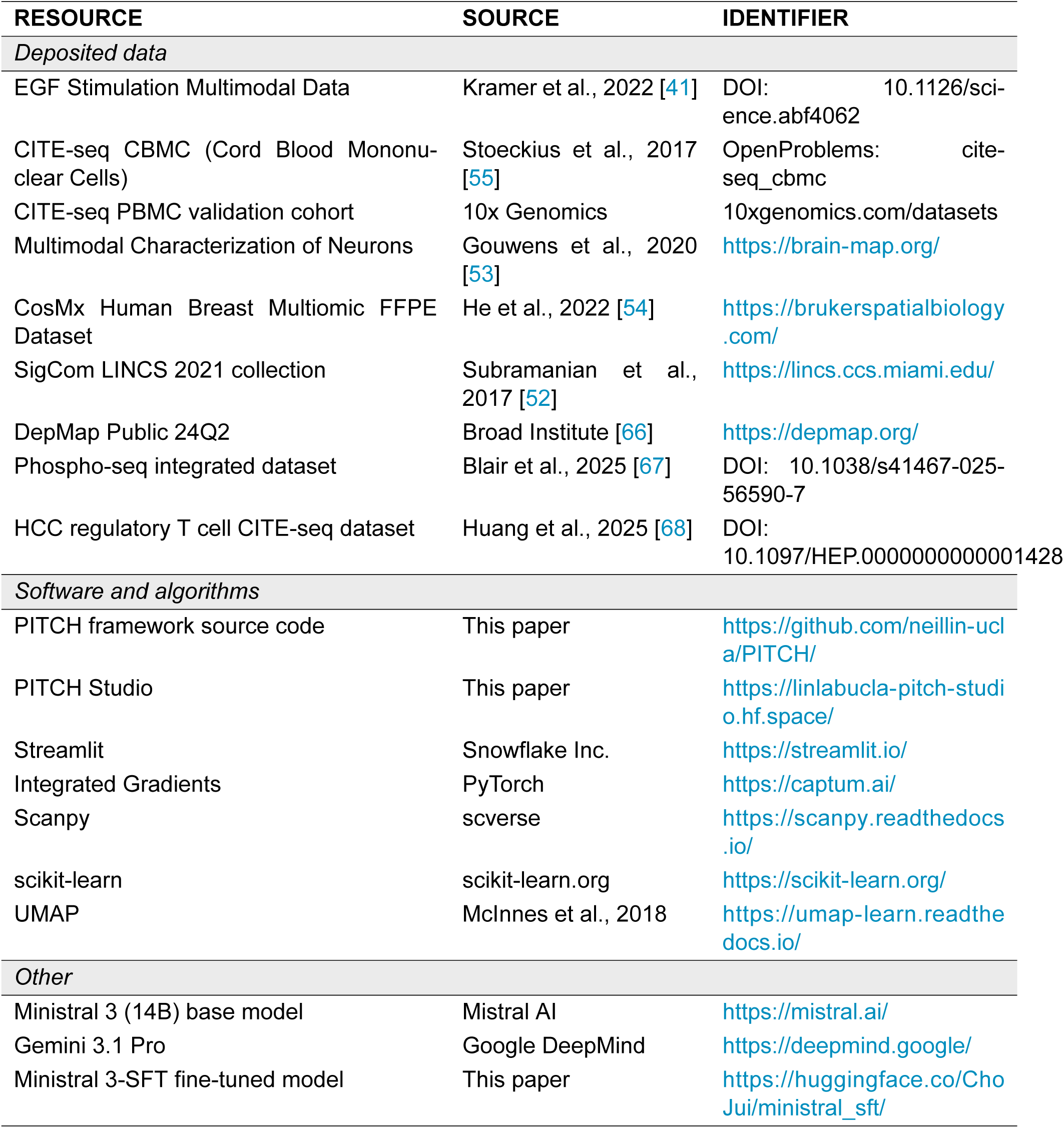

### METHOD DETAILS

#### Multimodal Data Compilation and Preprocessing Pipeline

To establish a biologically accurate zero-vector baseline for downstream IG extraction, raw multimodal-omics count matrices were uniformly processed through a core standardization pipeline. Across applicable datasets, low-quality cells were removed by filtering instances containing fewer than 200 gene counts or expressing less than 70% of total target genes. Retained cells then underwent sequencing depth normalization followed by z-score standardization, scaling the mean expression of each feature to zero with unit variance. Dimensionality reduction was subsequently applied prior to MLP training by isolating features with the 1,000 highest population variances, which were ranked and filtered by their Pearson correlation to designated target markers.

Specific datasets required targeted preprocessing steps. For the EGF stimulation multimodal dataset dataset [41], features exhibiting technical artifacts, including raw-to-log2 transform mismatches, repetitive features, and ambiguous background annotations, were excluded, reducing the feature space from 650 to 510 prior to log2-normalization and final z-scaling. The CosMx Human Breast Multiomic FFPE dataset [54] was acquired from Bruker Spatial Biology. Preprocessing for this spatial dataset involved computationally depleting infiltrating blood cells by thresholding PanCk and CD45 expression parameters. The Patch-seq dataset [53] was sourced from the Allen Institute for Brain Science, and the CITE-Seq Cord Blood Mononuclear Cell (CBMC) dataset was acquired from Stoeckius et al. [55]. For the CITE-seq PBMC validation cohort, raw transcriptomic counts were scaled via total-count normalization (10^4^ reads/cell) and log1p transformation, incorporating pre-computed Louvain annotations without supplementary feature pruning.

The SigCom LINCS 2021 collection [52] and the DepMap 24Q2 database [66] were both acquired from the Broad Institute. Preprocessing for the LINCS data required Characteristic-Direction coefficient vectors to be column-wise z-standardized, while the DepMap bulk expression data (TPM log1p) was filtered for the breast lineage and mapped to HGNC symbols. Finally, retrospective mechanism reconstructions utilizing the Phospho-seq [67] and hepatocellular carcinoma microenvironment [68] datasets were modeled on transcriptomic and surface protein modalities. To prevent target leakage during HCC modeling, regulatory T cells were isolated utilizing a strict FOXP3-independent marker panel. Fully reproducible processing scripts for all dataset iterations are provided in the Supplemental Information.

#### Multilayer Perceptron and EGF Encoder Architecture

We implemented a two-layer Multilayer Perceptron (MLP) to map high-dimensional cellular features to target biomarker abundance. To mitigate overfitting across all models, network bias terms were explicitly disabled and a dropout rate of 0.5 was applied. Consequently, the base transformation is defined mathematically as f(X) = *σ*(XW_0_)W_1_ where 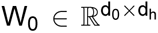 and 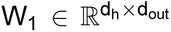 represent the learned weight matrices. Here, d_0_ denotes the input feature dimensionality, d_h_ is the hidden layer dimensionality, d_out_ is the output dimensionality, and *σ*(·) represents the Rectified Linear Unit (ReLU) activation function.

Model optimization was executed using the Adam optimizer. For the independent CITE-seq blood cell, Patch-seq single-neuron, phospho-seq and hepatocellular carcinoma microenvironment datasets, the hidden layer was configured with d_h_ = 256 nodes. These base models (d_out_ = 1) were trained for a maximum of 2,000 epochs utilizing a learning rate of 2 × 10^−4^ and a weight decay of 1 × 10^−4^. To accommodate the heightened complexity of modeling RNA-to-protein translation states within the CosMx spatial transcriptomics dataset, the hidden layer capacity was expanded to d_h_ = 512 nodes, maintaining all other base hyperparameters.

To capture context-dependent signaling dynamics in the EGF stimulation dataset, the base MLP was adapted into a condition-specific architecture. The network was modified to output a multi-dimensional latent vector (d_out_ = 256), which was subsequently evaluated against a learned EGF encoder embedding matrix, 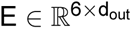, where 6 represents the categorical EGF stimulation conditions (0, 1, 6.25, 10, 25, and 100 ng/mL). During forward propagation, the specific stimulation condition for a given sample indexes the corresponding row vector from E. The final scalar prediction is computed via a dot product between the cell’s latent feature vector and this condition-specific weight vector. This design isolates condition-agnostic phenotypes within the shared MLP weights, while the encoder captures dose-specific nonlinear scaling. This specialized implementation utilized a hidden layer of d_h_ = 256 nodes and was trained for 40,000 epochs with a learning rate of 2 × 10^−4^ and a weight decay of 2 × 10^−3^ (Table S4).

#### Benchmarking of Predictive Regression Models

To evaluate the predictive performance of the MLP architecture, we benchmarked it against a standard multilinear baseline (ordinary least squares) and five representative nonlinear machine learning models. The nonlinear baselines included a distance-weighted k-Nearest Neighbors regressor (k = 20), an RBF kernel ridge regressor (accelerated via Nyström approximation), and three tree-based ensembles: Random Forest, Histogram-based Gradient Boosting, and LightGBM. Model performance was assessed using 5-fold cross-validation with randomized data shuffling. Input features for linear, distance-based, and kernel-based estimators were standardized within the cross-validation pipeline, whereas tree-based algorithms operated on unscaled features. Regularized baseline models optimized their penalty hyperparameters via inner cross-validation to prevent artifactual underperformance. Predictive accuracy across the models was quantified using the Root Mean Square Error to measure residual error in the original biological units, which was subsequently used to calculate proportional residual reduction relative to the linear baseline. To ensure generalizability, this benchmarking protocol was executed independently across 12 distinct signaling responses (e.g., pRSK, pERK, pS6, and pEGFR) (Table S5).

#### Integrated Gradients Calculation and Axiomatic Properties

Let f : ℝ^d^ → ℝ represent the trained neural network, mapping an input vector x = [x_1_, x_2_, *…*, x_d_]^T^ ∈ ℝ^d^ relative to a reference baseline x*^′^* ∈ ℝ^d^. IG quantifies the precise contribution of each feature to the overall change in model output, transitioning from the baseline state f(x*^′^*) to the final prediction f(x), by integrating the mathematical gradients along a continuous straight-line path from x*^′^* to x.

The attribution for the i-th feature, denoted as G_i_(x), is defined as 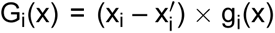 where g_i_(x) represents the integrated gradient:

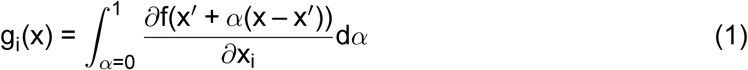

For linear functions, this attribution process simplifies, as g_i_(x) reduces to a constant equivalent to the learned coefficient of x_i_.

A fundamental mathematical property of IG is the completeness axiom, which dictates that the sum of all feature attributions must exactly equal the difference between the model’s output at x and the output at the baseline x*^′^*. For any function differentiable almost everywhere, this relationship is expressed as 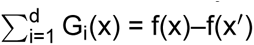. This axiom guarantees that IG fully accounts for the contribution of all input features, providing a complete decomposition of the model’s prediction. Because the input features undergo z-score standardization during preprocessing, establishing the baseline x*^′^* as the zero vector represents the average population expression state. Under this condition, the completeness relationship reformats to 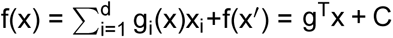 where g = [g_1_(x), g_2_(x), *…*, g_d_(x)]^T^ and the constant C = f(x*^′^*).

Finally, IG satisfies the axiom of Implementation Invariance. If two neural networks are functionally equivalent, producing identical outputs for all inputs, they yield identical attributions regardless of variations in their internal architectures, parameters, or hidden layers. This property ensures that the extracted attributions reflect the input-output dynamics of the biological data rather than arbitrary algorithmic implementations.

#### Decision Tree Logic Extraction

To convert non-linear IG attributions into logical rules, we implemented a decision tree classifier utilizing the CART (Classification and Regression Trees) algorithm [88]. To establish discrete classification targets, we isolated the distributional tails of the IG scores for demonstrations: cells exhibiting the top 10% of attribution scores were designated as the positive condition, representing strong phenotypic feature dependence, while the background population was designated as the negative condition.

The classifier was trained on the standardized cellular feature expression profiles to predict these discrete IG classes. At each internal node, the algorithm recursively partitions the feature space by evaluating all possible thresholds to minimize the Gini impurity, defined mathematically as: H(Q_m_) = Σ_k_ p_mk_(1 – p_mk_) where p_mk_ represents the proportion of observations belonging to class k within node m. To ensure that the extracted Boolean logic remained interpretable and to prevent computational overfitting, the tree architecture was regularized. Maximum tree depth was constrained to 4, and strict minimum sample thresholds were enforced for all internal node splits. Furthermore, to counteract the class imbalance introduced by isolating the attribution tail, a class weight multiplier of 4 was applied to the positive minority class during training. This penalty scales the Gini impurity reduction criterion, forcing the algorithm to prioritize the accurate capture of the rare, highly attributed cellular phenotypes over majority-class background accuracy.

#### Browser-Based Pipeline Deployment and Software Architecture

The PITCH Studio framework was deployed as a browser-based application built in Python using the Streamlit library. To ensure portability, the architecture operates without a persistent database, leveraging Streamlit’s session state for the in-memory management of high-dimensional matrices, tensors, and intermediate algorithmic artifacts. The frontend interfaces with a core analytical backend, where UI state transitions programmatically trigger MLP optimization (PyTorch), IG extraction (Captum), and decision tree induction (scikit-learn). Visualizations of tree topologies and feature distributions are dynamically rendered using Matplotlib, Plotly, and NetworkX. Finally, automated hypothesis generation is facilitated by a RESTful API module, which packages the extracted Boolean logic into structured JSON payloads and securely queries the Gemini API to retrieve and render asynchronous mechanistic interpretations.

#### Performance Evaluation of Large Language Models

To benchmark the mechanistic interpretability of both our fine-tuned and commercial LLMs, we implemented an automated “LLM-as-a-judge” evaluation framework utilizing Gemini 3.1 Pro as the designated teacher evaluator. Model performance was assessed using a hold-out test set comprising 200 unseen synthetic logic prompts, which served as functionally validated ground-truth targets. To quantify interpretative quality, candidate model generations were evaluated against a structured five-metric rubric. The teacher model scored each response based on biological plausibility, logic rigor, literature grounding, feature comprehensiveness, and mechanistic specificity. The mean scores across the 200-prompt cohort were aggregated to establish comparative performance baselines.

Simultaneously, we implemented a penalization protocol to quantify the algorithmic failure rate of the models. Model generations were screened and flagged for five distinct modes of reasoning failure: pathway hallucination, context mismatch, causal overreach, logic failure, and vagueness (Table S6). Finally, the overall accuracy of the LLM-generated hypotheses was benchmarked directly against the foundational ground-truth rules. To capture the nuance of biological interpretation, response quality was not treated as a binary variable but rather stratified into four distinct qualitative tiers: a perfect match, highly relevant, loosely relevant, or entirely irrelevant to the fundamental ground-truth mechanism. My replacement: To evaluate the LLM’s interpretability independently of biases introduced by tree synthesis, we filtered out tree prompts that failed to reconstruct the ground-truth mechanism.

#### Language Model Fine-Tuning and Hypothesis Generation

To enable local, privacy-preserving mechanistic interpretation, we performed knowledge distillation from Gemini 3.1 Pro into the open-weight Ministral 3 (14B parameter) architecture via full Supervised Fine-Tuning (SFT). The training corpus comprised 30,000 synthetic decision tree logic prompts spanning diverse biological domains. Gemini 3.1 Pro was queried to generate high-fidelity biological interpretations alongside explicit Chain-of-Thought (CoT) reasoning for each prompt. This paired corpus of Boolean boundary conditions and CoT-augmented mechanistic narratives was subsequently utilized to optimize the custom Ministral 3-SFT model.

For applied hypothesis generation across the empirical cohorts (Patch-seq, CITE-seq, and CosMx spatial transcriptomics datasets), the algorithmically extracted Boolean rules were formulated into structured text prompts. Mechanistic inference was conducted using Gemini 3.1 Pro, with the sampling temperature explicitly set to 1.0. This parameterization was selected to maximize generative diversity and facilitate broad, cross-disciplinary literature synthesis when translating complex numerical attributions into testable biological hypotheses.

#### Empirical Validation of LLM-Generated Hypotheses

To validate actin-dependent signaling predictions, transcriptional responses to pharmacological actin depolymerization (latrunculin-B, cytochalasin-D, and cytochalasin-B) and EGF stimulation (positive controls) were queried from the SigCom LINCS 2021 collection. Characteristic-Direction (CD) coefficient vectors spanning 83 signatures across epithelial lineages (MCF10A, A549, MCF7, and HA1E) were column-wise z-standardized. Regulatory dependencies within a targeted 7-gene MAPK module and a 13-gene YAP/TAZ module were subsequently evaluated against the genome-wide background distribution utilizing two-sample Kolmogorov–Smirnov tests.

To assess nominated genetic dependencies, Chronos CRISPR gene-effect scores and bulk transcriptomic profiles (TPM log_1p_) were obtained from the DepMap 24Q2 database. Data were restricted exclusively to the breast lineage and mapped to HGNC symbols. For essentiality benchmarking across the 51 identified breast cancer models, per-gene mean Chronos scores and 95% confidence intervals (CIs) were computed; statistical support for differential genetic essentiality was predefined as non-overlapping CIs (e.g., between *PTMA* and *RPLP0*). For mitotic co-expression modeling, uncorrected pairwise Pearson correlations were computed among target markers, with composite signatures derived as the per-model mean of their column-wise z-scores. All statistical analyses were executed in Python (pandas, scipy).

To empirically test LLM-nominated immune mechanisms, the independent PBMC validation cohort (n = 2,638) was scaled via total-count normalization (10^4^ reads/cell) and log_1+x_ transformation. Louvain-annotated cytotoxic lineage cells (e.q., NK cells and CD8^+^ T cells) were used as the reference population for threshold estimation. To accommodate zero-inflated droplet distributions, PITCH-zone boundaries were approximated as cytometric-equivalent inequalities based on the 25th (P_25_) and 75th (P_75_) expression percentiles of gate-defining lineage markers. Mature lineage gating makers are *KLRD1*,> P_75_, *NKG7*,> P_75_, *GZMB*,> P_75_, and *CST3*,≤ P_25_; Primed T cells required *NKG7*,> P_75_, *GZMB*,≤ P_25_, and *S100A12*,≤ P_25_. Remaining cytotoxic lineage cells outside the nominated PITCH zones were used as the biologically matched baseline. Cytotoxic activity was quantified using a nine-gene module score (*GZMB*, *GZMA*, *GZMK*, *PRF1*, *KLRD1*, *NKG7*, *GNLY*, *CCL5*, *FGFBP2*) computed with a Seurat-style expression-bin control method. Directional mechanistic hypotheses generated by the LLM, including selective *GNLY* priming prior to coordinated *GZMB*-associated cytotoxic maturation, were evaluated using stratified Mann-Whitney U tests and Spearman correlation analysis. To isolate lineage-specific effects from baseline confounding, *GNLY* enrichment was tested against the matched cytotoxic baseline, with effect sizes reported as rank-biserial correlations (rbc) and significance defined as p < 0.05.

### QUANTIFICATION AND STATISTICAL ANALYSIS

#### Training Performance Evaluation

The processed dataset was randomly partitioned into training (80%) and testing (20%) data splits. Following model training, the model was applied to predict the target values using the testing data. The Pearson correlation between the predicted target values and the measured target values was then calculated to evaluate model training performance.

#### Gini Impurity and Precision

To iteratively partition the data and isolate target cellular states, the decision tree algorithm optimizes for node purity by minimizing the Gini impurity at each split. Gini impurity is calculated as 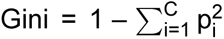 where C represents the total number of classes (C = 2 for our binary classification scheme), and p_i_ is the probability of a sample belonging to class i within a given node. A perfectly pure leaf, where the probability of selecting the target class is 1, yields a Gini impurity of 0. To quantitatively evaluate the reliability of terminal leaves, we calculated the precision for each defined rule 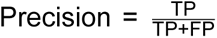 where True Positives (TP) denotes the number of correctly classified target samples within the leaf, and False Positives (FP) represent the number of non-target samples captured within the same leaf.

