## Supplemental Figures for "Synthesizing Mechanistic Hypotheses from Single-Cell Omics via Discretized Feature Attribution and Empirical Language Model Grounding"

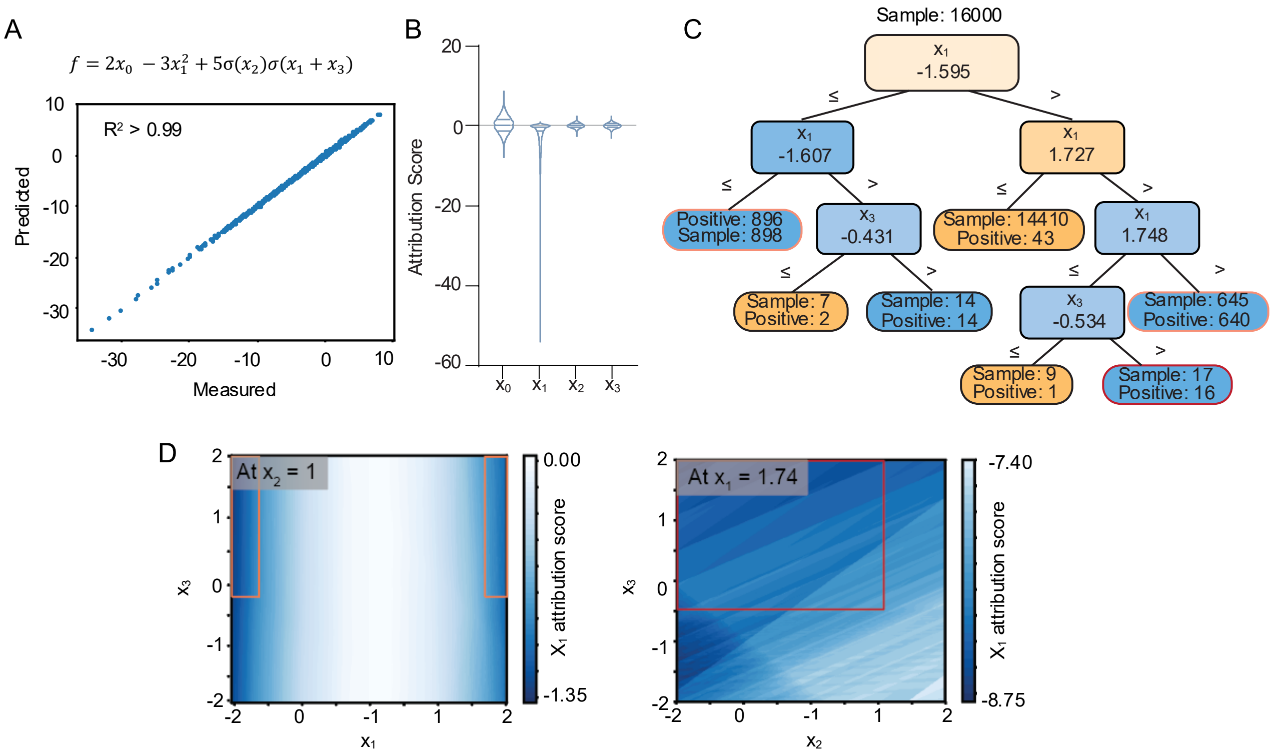


**Figure S1. Validation of the computational framework capability using synthetic dataset.** (A) A synthetic nonlinear equation, designed to mimic complex biological regulation, is utilized to evaluate the predictive performance of the MLP. Applying MLP achieves accurate predictions of $f$ ($R^2 > 0.99$), demonstrating its capacity for modeling nonlinear dynamics. (B) Feature attribution via IG reveals that the dominant variable, $x_1$, exerts a significant negative contribution to $f$. (C) DT classifier delineates a rule-defined subset corresponding to the bottom 10% $x_1$ IG scores. High-precision terminal nodes are highlighted with orange-frame zones (sample = 645, positive = 640; sample = 898 and positive = 896) and a red-frame zone (sample = 17, positive = 16), differentiated by their dependency on the $x_3$ splitting criterion. (D) Heatmaps visualize the distributions of the $x_1$ attribution scores corresponding to the orange and red zones. Under the condition $x_2 = 1$, the $x_1$ attribution score largely independent of $x_3$. However, when constrained to $x_1 = 1.74$, the $x_1$ attribution score relies more on $x_3$ than $x_2$.

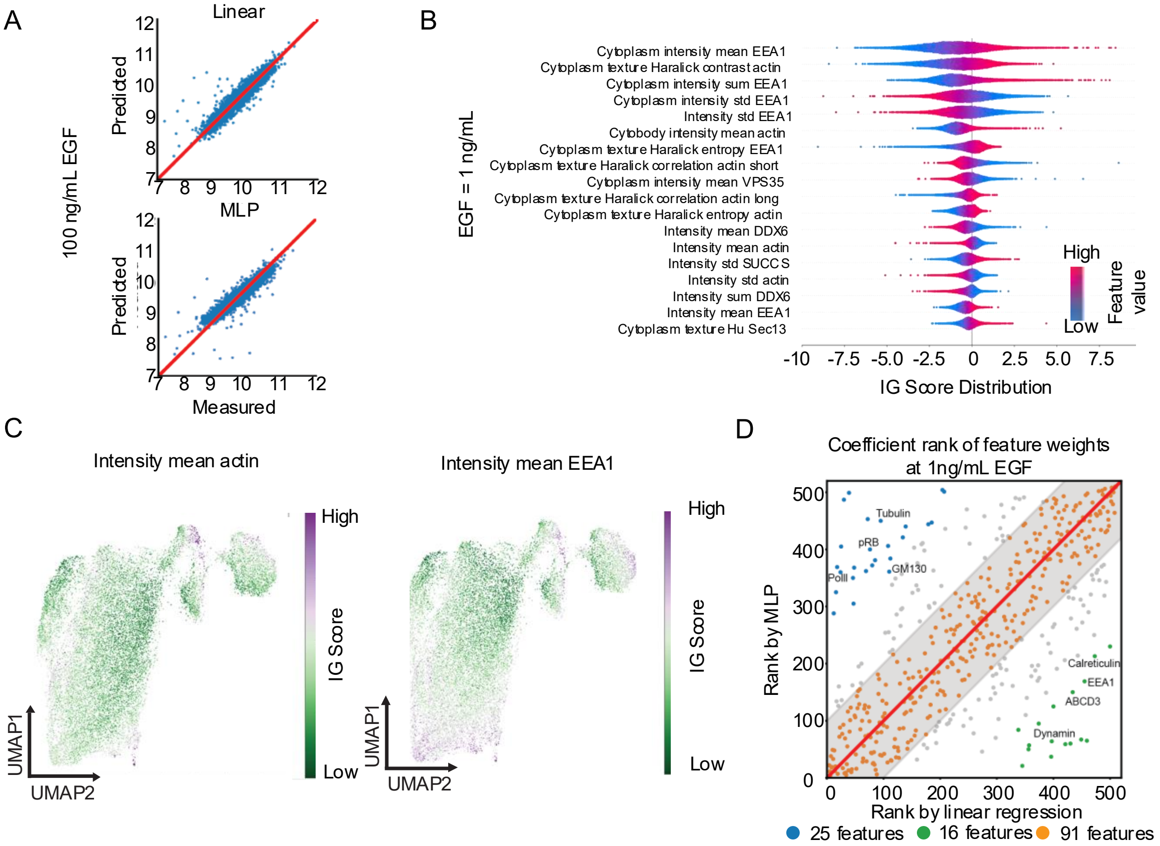


**Figure S2. MLP and IG analysis at low EGF concentration with high-concentration performance baselines.** (A) At saturating 100 ng/mL EGF, both linear regression and MLP show accurate predictions of pERK. (B) Ranking the IG scores reveals that EEA1 and Actin are the primary predictors, both positively correlated with pERK activity at 1 ng/mL EGF. (C) The projections of Actin and EEA1 IG scores on the UAMP show a closed distribution. (D) Scatter plot of the feature weights ranked by linear regression against by MLP at 1 ng/mL EGF. Green dots highlight components of the receptor internalization machinery (e.g., EEA1, Dynamin, Calreticulin) whose complex dynamics are prioritized by the MLP. Conversely, blue dots represent distributed cell cycle markers (e.g., Pol II, pRB, Tubulin) highlighted by linear regression. Orange dots denote similarly ranked features.

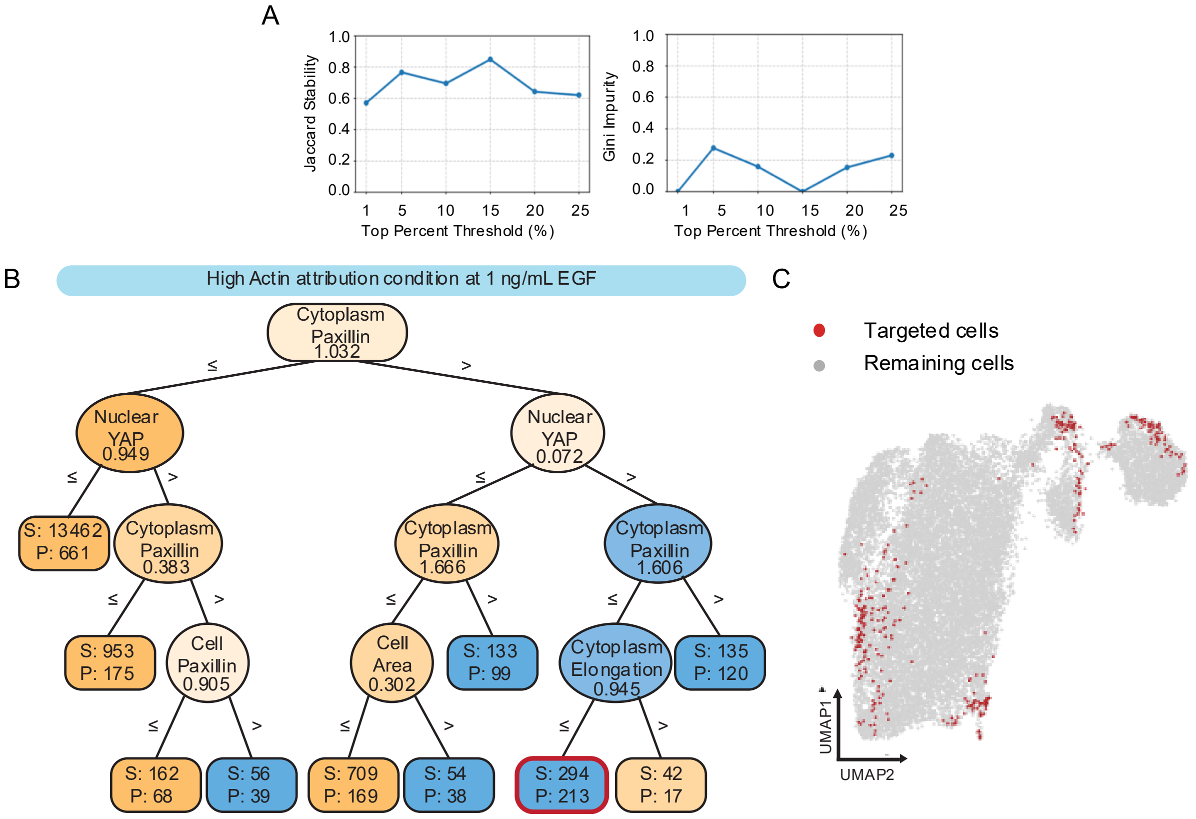


**Figure S3. Characterization of target cell threshold in DT classification.** (A) Characterization of the IG score threshold for DT classification at 10 ng/mL EGF. Concurrently, the Jaccard stability for the top 5 features exceeds 0.7 between the 5% and 15% thresholds, indicating high reproducibility within this range. Low Gini scores occur at thresholds between 10% and 20%, where the target cell population comprises over 80% of the total cells. (B) Using the same 22 mechanobiology features, the DT identifies a high-precision subpopulation at 1 ng/mL EGF marked by elevated cytoplasmic paxillin and nuclear YAP, with low cytoplasmic elongation. S: total sample size; P: positive sample size. (C) UMAP projection highlighting the distribution of cells from the red-framed leaf in (B), demonstrating alignment with high actin IG scores (Figure S2C, left).

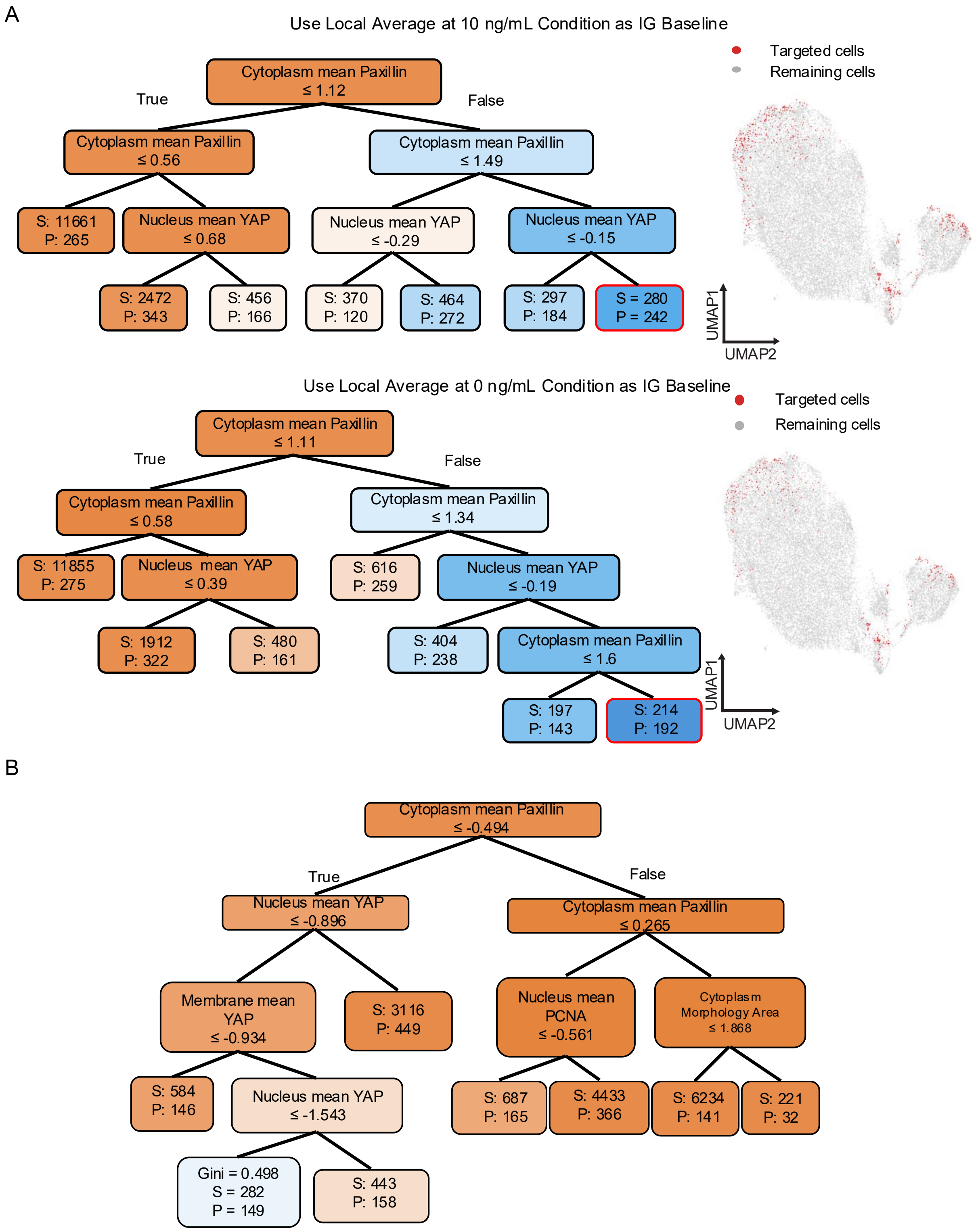


**Figure S4. Control analyses confirm the robustness and specificity of extracted biological rules.** (A) DT classifications robustly utilize cytoplasmic paxillin and nuclear YAP to identify cells with high actin attribution at 10 ng/mL, showing consistency between the local baseline (top) and biological neutral baseline (bottom) used for IG calculation. UMAP projections of the identified subpopulations (red-framed boxes) remain consistent across both baselines. S: total sample size; P: positive sample size. (B) When applying the same 22 mechanobiology features to classify conditions where actin negatively contributes to pERK activation, the DT failed to isolate this state (maximum precision = 53%).

HIGHLIGHT ZONE 1: The Mechanotransduction-Scaffold Axis (Node 28)
Logic: Cytoplasm Paxillin > 1.342 AND Nuclear YAP > -0.236 AND Cytobody Paxillin > 1.597
Performance: True Positives = 248, Precision = 89.2%
This is the most biologically robust zone, capturing the highest number of true positives with near-maximum precision.

Biological Rationale:
This node describes a cellular state characterized by highly abundant and widely distributed Paxillin (both cytoplasmic and whole-cytobody) coupled with non-negative/high nuclear YAP. Biologically, this represents a state of high cytoskeletal tension and mature focal adhesion assembly. When breast epithelial cells are stimulated with EGF, the EGFR pathway (leading to pERK) does not operate in a vacuum; it heavily cross-talks with integrin signaling. Paxillin is a core component of focal adhesions. High tension (indicated by nuclear YAP, a known mechanosensor) drives the polymerization of globular actin (G-actin) into filamentous actin (F-actin) stress fibers. In this specific high-tension state, the Actin cytoskeleton acts as an essential spatial scaffold. It physically clusters EGFRs at the membrane and anchors the RAS/RAF/MEK/ERK signaling cascade via actin-binding scaffolding proteins (e.g., IQGAP1). Therefore, in cells with mature focal adhesions and active YAP, the state of the Actin cytoskeleton becomes the ultimate bottleneck/predictor for how efficiently EGF translates into pERK.

Findings Directly Confirming Literature:
Integrin-EGFR Crosstalk: The reliance on Paxillin to contextualize the Actin-pERK relationship confirms that integrin-mediated adhesion is required for optimal growth factor signaling. Actin as a Signaling Scaffold: The high attribution of Actin to pERK in this state aligns with the known role of F-actin in scaffolding the MAPK cascade to prevent signal dissipation. YAP Mechanosensing: The presence of nuclear YAP alongside high Paxillin confirms the canonical Hippo pathway mechanotransduction model, where cytoskeletal tension drives YAP nuclear translocation.

Hypothesis:
The strong predictive power of Actin on pERK in this state is driven by a YAP-mediated positive feedback loop. We hypothesize that nuclear YAP drives the transcription of specific actin-crosslinking proteins (e.g., Filamin A) or EGFR ligands (e.g., Amphiregulin), which in turn stabilize the Actin-EGFR-MEK complex. Experimental Test: Knockdown YAP in these cells, stimulate with EGF, and observe if the spatial co-localization of Actin and pERK (via immunofluorescence) is decoupled, and if Actin loses its predictive power for pERK.

Supporting Publications:
Dupont et al. (2011), Nature: Established that YAP is a nuclear relay of mechanical signals exerted by extracellular matrix rigidity and cell shape, requiring the actin cytoskeleton.Miyamoto et al. (1996), Journal of Cell Biology: Demonstrated the direct crosstalk between integrin signaling (Paxillin) and growth factor receptors (EGFR/MAPK) via cytoskeletal scaffolding. Bisson et al. (2011), Cell Research: Highlighted the role of IQGAP1 and actin in scaffolding the MEK/ERK pathway

**Figure S5. Full tree interpretation generated by the LLM for 10 ng/mL EGF perturbation.** The LLM identifies the condition with high actin attribution and suggests rationale, testable hypothesis and supporting publications from the decision tree logic. It proposes that actin cytoskeleton scaffold maintained by mechanotransduction (high nuclear YAP) clusters EGFRs and thus promotes the cross-talked RAS/RAF/MEK/ERK signaling cascade.

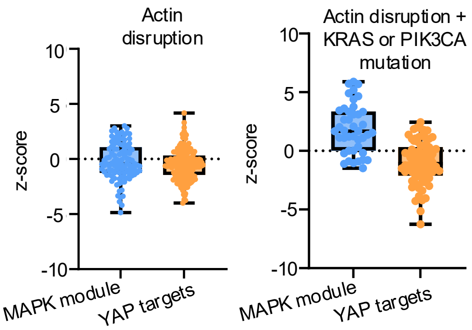


**Figure S6. Characterization and downstream functional analysis of Actin-driven subpopulations.** Pharmacological actin depolymerization in epithelial cells suppresses both MAPK activation and YAP/TAZ transcriptional targets (n = 15 signatures, top). Constitutively active KRAS or PIK3CA mutations restore MAPK output relative to actin depolymerization alone (n = 8 signatures, bottom; $p < 10^{-8}$), but fail to rescue YAP/TAZ target expression.

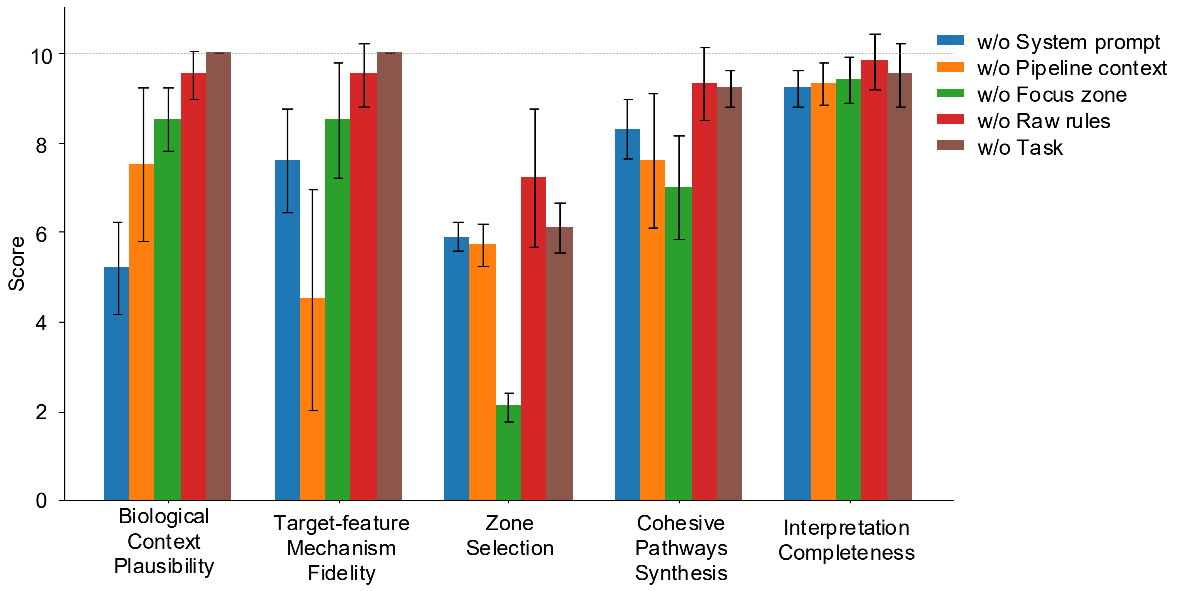


**Figure S7. Ablation testing of the structured prompt within the PITCH framework.** Evaluation across five performance metrics (scored 1–10) characterizes the relative contribution of individual prompt components to LLM performance. Removal of the system prompt significantly degrades *Biological Context Plausibility by about* 5 point, likely due to the loss of background framing and alignment constraints. Omitting the pipeline context reduces *Target-Feature Mechanism Fidelity*, indicating that the model requires structured analysis guidance to interpret contextual dependencies. Excluding selected focus zone candidates decreases *Zone Selection* accuracy (by 50%) and impairs *Cohesive Pathways Synthesis*. Conversely, removing raw rules has a negligible impact, emphasizing the hierarchical importance of prompt architecture. Because core task instructions are reinforced throughout other prompt modules, excluding the explicit task section does not significantly alter LLM performance.

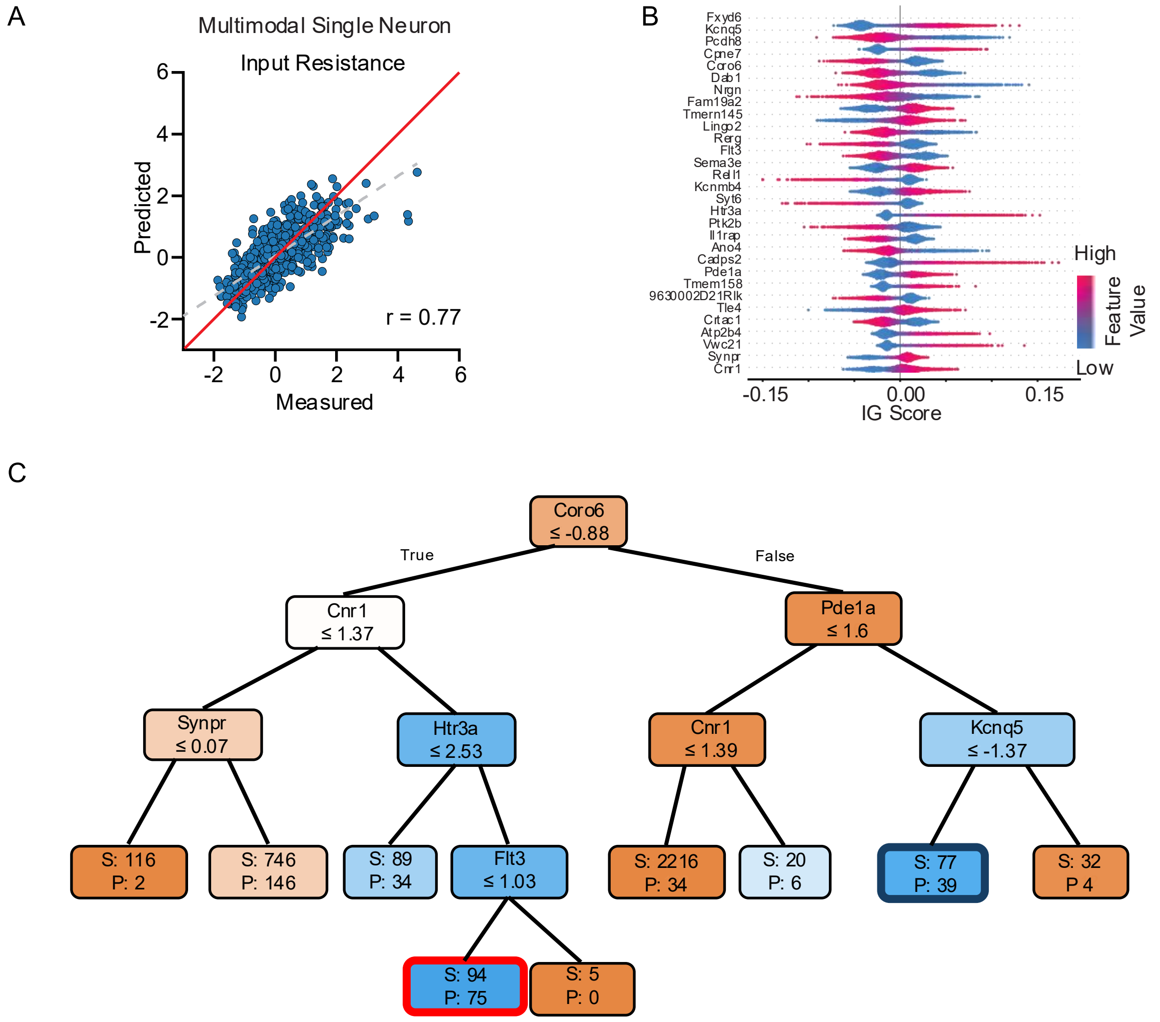


**Figure S8. PITCH identifies the specific cellular context involved in regulating input resistance.** (A) Scatter plot of measured versus predicted input resistance, demonstrating that the MLP model accurately predicts input resistance (Pearson's r = 0.77, gray dash line; low residual error, red identity line) from single-neuron RNA profiles. (B) A beeswarm plot of IG scores reveals that ion channel and pump genes are the dominant predictors of input resistance, with active-pump regulator *Fxyd6* ranking as the top feature. (C) The DT extracts conditional rules identifying subpopulations where *Fxyd6* most strongly associates with input resistance: a low-*Coro6*, high-*Cnr1*, high-*Htr3a*, and low-*Flt3* subpopulation (condition 1, red frame), alongside a high-*Coro6*, high-*Pde1a*, and low-*Kcnq5* subpopulation (condition 2, blue frame).

PRIMARY ZONE: Node 23 (The CGE-Derived CCK+ Interneuron State)
Logic: Coro6 <= -0.877 AND Cnr1 > 1.370 AND Htr3a > 2.532 AND Flt3 <= 1.030
Performance: True Positives: 75 | Precision: 79.8% | Gini: 0.1120.

Biological Rationale:
This node perfectly describes a well-known subpopulation of cortical interneurons. Htr3a (Serotonin receptor 3A) is the canonical marker for all CGE-derived interneurons. Within the Htr3a+ population, high expression of Cnr1 (Cannabinoid Receptor Type 1 / CB1R) specifically identifies Cholecystokinin (CCK+) basket cells.In this specific CGE-derived (Htr3a+/Cnr1+) developing neuron population, Fxyd6 becomes a critical predictor of input resistance. Because developing neurons often lack the full suite of mature "leak" potassium channels, the electrogenic nature of the Na+/K+ ATPase pump (which Fxyd6 modulates) becomes a primary driver of resting membrane potential and, consequently, resting membrane conductance (input resistance).

Findings Confirming Literature Knowledge:
Marker Co-expression: The co-expression of Htr3a and Cnr1 accurately identifies CGE-derived CCK+ interneurons, confirming the model has isolated a biologically valid cell type. Fxyd6 Function: Fxyd6 is known as phosphohippolin, a tissue-specific modulator of Na+/K+ ATPase. Modulating this pump directly alters ion gradients and electrogenic currents, which are known to influence input resistance, especially in compact, developing neurons with high baseline input resistance.

New Testable Mechanisms & Hypotheses:
Hypothesis 1 (Receptor-Pump Coupling): We hypothesize that in developing CGE-derived interneurons, 5-HT3R (Htr3a) and CB1R (Cnr1) signaling cascades functionally converge to regulate Fxyd6 activity or its interaction with the Na+/K+ ATPase. Testable Mechanism: Agonism of CB1R in developing CGE neurons alters input resistance only when Fxyd6 is present, suggesting Fxyd6 acts as a downstream effector of endocannabinoid signaling to regulate membrane excitability during development. Hypothesis 2 (Developmental Excitability Switch): Flt3 (low in this node) is often associated with early progenitor proliferation. Its absence, combined with high Cnr1, suggests a post-mitotic, migrating/maturing state.

Testable Mechanism: Fxyd6-mediated modulation of input resistance is a transient developmental requirement for CGE neurons to integrate into early cortical circuits before mature potassium leak channels (like TASK channels) take over resting conductance.

SECONDARY ZONE: Node 15 (The M-Current Deficient / cAMP-Active State)
Logic: Coro6 > -0.877 AND Pde1a > 1.599 AND Kcnq5 <= -1.366
Performance: True Positives: 39 | Precision: 50.6% | Gini: 0.3150.

Biological Rationale: This node isolates a state defined by low Kcnq5 and high Pde1a. Kcnq5 encodes the Kv7.5 potassium channel, a primary subunit of the M-current, which is a dominant regulator of resting membrane potential and input resistance (low Kcnq5 = fewer open channels = higher input resistance). Pde1a is a Calcium/Calmodulin-dependent phosphodiesterase that hydrolyzes cAMP/cGMP. When M-current (Kcnq5) is low, the membrane is "tight." In a tight membrane, the relative contribution of the Na+/K+ ATPase pump to total membrane conductance is mathematically magnified. Therefore, Fxyd6 (modulating the pump) becomes a highly sensitive, dominant predictor of input resistance. Furthermore, high Pde1a suggests active regulation of cyclic nucleotides, which are known to phosphorylate and modulate both ion channels and Na+/K+ pumps.

Findings Confirming Literature Knowledge: Kcnq5 and Input Resistance: It is well-established that KCNQ/Kv7 channels dictate neuronal input resistance. A reduction in these channels increases input resistance.PDE1A and Excitability: Phosphodiesterases regulate intracellular cAMP/cGMP, which in turn regulate Protein Kinase A/G (PKA/PKG). PKA is known to phosphorylate the Na+/K+ ATPase, altering its affinity and activity.

New Testable Mechanisms & Hypotheses:
Hypothesis 1 (Biophysical Synergy): We hypothesize a compensatory biophysical relationship where Fxyd6-regulated Na+/K+ ATPase activity becomes the primary determinant of input resistance specifically when Kv7.5 (Kcnq5) M-currents are developmentally delayed or suppressed. Testable Mechanism: Pharmacological blockade of Kv7 channels (e.g., with XE991) in MGE/CGE neurons will exponentially increase the effect size of Fxyd6 knockout on input resistance. Hypothesis 2 (cAMP-Dependent Pump Modulation): High Pde1a indicates rapid turnover of cAMP. We hypothesize that Fxyd6 requires a specific cyclic nucleotide environment (regulated by Pde1a) to effectively couple with the Na+/K+ ATPase.

Testable Mechanism:
Inhibition of PDE1A will alter cAMP levels, leading to hyper-phosphorylation of the Na+/K+ ATPase, thereby decoupling Fxyd6 from the pump and abolishing Fxyd6's predictive power over input resistance.

**Figure S9. Full tree interpretation by the LLM for input resistance regulation.** The model identifies cellular conditions with high Fxyd6 association and proposes biological rationales, testable hypotheses, and supporting literature based on the decision tree logic. It highlights two subpopulation-specific regulatory mechanisms, suggesting that reduced M-currents shift input resistance dependency toward active transport to maintain homeostasis.

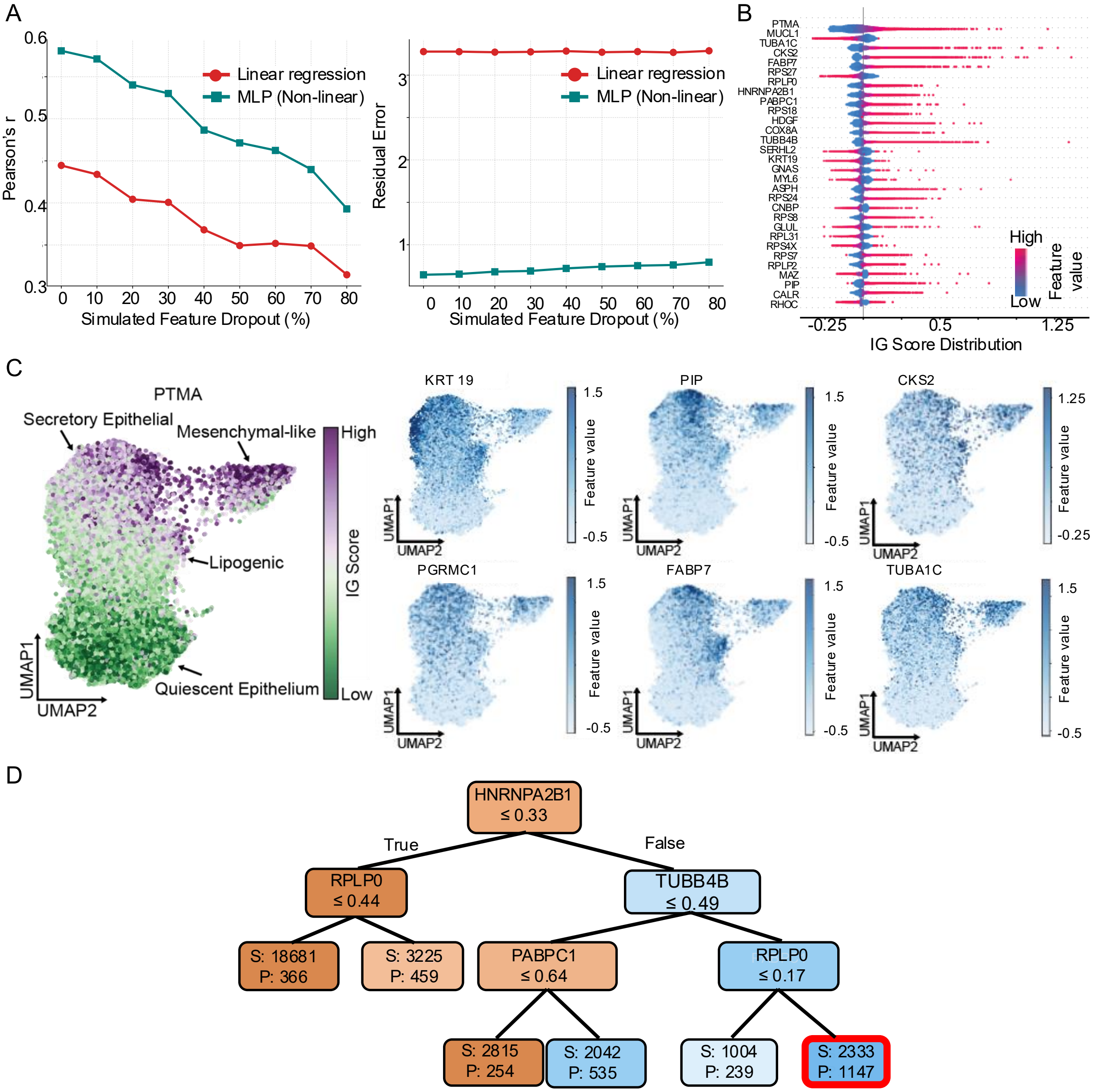


**Figure S10. PITCH reveals the regulatory logic of breast cancer proliferation.** (A) Using intact RNA features, linear regression showed limited accuracy for predicting single-cell Ki-67 abundance (Pearson's r = 0.44; residual error ≈ 3.3), whereas the MLP improved prediction performance (Pearson's r = 0.59) and reduced residual error by approximately 5.5-fold. Under simulated feature dropout, the MLP maintains higher Pearson's r and approximately 4–5-fold lower residual error than the linear baseline across dropout levels; even at 70% feature loss, its accuracy (r = 0.44) remained comparable to the intact-feature linear model. (B) Ranking feature importance by IG scores for the MLP-based Ki-67 prediction, identifying chromatin-associated regulator *PTMA* as the primary predictor. (C) UMAP projection of *PTMA* IG score, demonstrates that its attribution score increases along a trajectory from secretory epithelial cells toward mesenchymal-like cells. *KRT19* and *PGRMC1* mark secretory epithelial identity, whereas *TUBA1C* and *CKS2* are enriched in the mesenchymal-like population. (D) The DT delineates a Ki-67-associated mitotic subpopulation by high-*HNRNPA2B1*, high-*TUBB4B* and high-*RPLP0.*

Based on the provided decision tree logic and the biological context of breast cancer, we can decode the regulatory environment that maximizes the predictive power of PTMA (Prothymosin alpha) on Ki-67 (MKI67).
PTMA is a known chromatin-remodeling protein and transcription regulator that prevents apoptosis and drives cell proliferation. Ki-67 is the gold-standard biomarker for cellular proliferation, expressed during active cell cycle phases (G1, S, G2, M). The decision tree reveals that PTMA does not drive Ki-67 in isolation; its predictive power is highly dependent on specific RNA processing, translational machinery, and cytoskeletal networks.

ZONE 1 ANALYSIS: The Primary Proliferation & Translation Axis (Node 14)
Logic: HNRNPA2B1 > 0.333 AND TUBB4B > 0.489 AND RPLP0 > 0.169
Performance: True Positives: 1147 | Precision: 51.4% (Most biologically robust zone)

Biological Rationale:
This node represents a highly active, dividing breast cancer cell state. For PTMA to strongly predict Ki-67, the cell must have an active RNA splicing/transport mechanism (high HNRNPA2B1), active translational machinery (high RPLP0), and active mitotic spindle formation (high TUBB4B). PTMA unwinds chromatin to facilitate the transcription of proliferation-associated genes. However, this transcriptional push only results in Ki-67 expression (and actual proliferation) if the downstream machinery can process the mRNA (HNRNPA2B1), translate it into protein (RPLP0), and physically execute cell division (TUBB4B).

Findings Directly Confirming Literature Knowledge:
PTMA and Ki-67 Coupling: Literature confirms that PTMA overexpression correlates with high Ki-67 indices and poor prognosis in breast cancer. Both are hallmarks of the S/G2/M phases of the cell cycle. HNRNPA2B1 as an Oncogene: HNRNPA2B1 is a known driver of breast cancer progression. It regulates alternative splicing and mRNA stabilization of oncogenes. Its high expression is required to process the massive transcriptional output initiated by chromatin remodelers like PTMA. Ribosomal and Cytoskeletal Demand: High expression of RPLP0 (ribosomal protein) and TUBB4B (tubulin) reflects the immense biosynthetic and structural demands of a proliferating cell preparing for mitosis.

New Testable Mechanisms & Hypotheses Hypothesis:
PTMA-driven upregulation of Ki-67 is post-transcriptionally gated by HNRNPA2B1-mediated mRNA processing and RPLP0-dependent translation. Testable Mechanism: While PTMA may increase the transcription of Ki-67 and other cell-cycle regulators, the actual protein expression of Ki-67 requires HNRNPA2B1 to stabilize or splice the transcripts. Proposed Experiment: Perform an in vitro knockdown of HNRNPA2B1 in high-PTMA breast cancer cell lines (e.g., MCF-7 or MDA-MB-231). We hypothesize that despite high PTMA levels, Ki-67 protein levels will plummet because the mRNA transcripts generated by PTMA's chromatin remodeling fail to be processed and translated, effectively uncoupling PTMA from Ki-67.

Supporting Literature References:
PTMA & Proliferation: Magdalena, C., et al. (2000). "Prothymosin alpha expression correlates with node-positive breast cancer and high proliferation index (Ki-67)." British Journal of Cancer. HNRNPA2B1 in Breast Cancer: Hu, Y., et al. (2017). “HNRNPA2B1 promotes breast cancer cell proliferation and metastasis..." Cancer Cell International. PABPC1 & mRNA Stability: Zhang, H., et al. (2021). "PABPC1 interacts with RNA to promote translation and proliferation in breast cancer." Oncogene. TUBA1C as a Biomarker: Li, Y., et al. (2022). "TUBA1C is a prognostic marker and correlated with immune infiltration in breast cancer." Frontiers in Oncology. Ribosomal Proteins (RPLP0) in Cancer: Artero-Castro, A., et al. (2009). "Expression of the ribosomal proteins RPLP0 and RPLP1 is upregulated in breast cancer." Molecular Cancer.

**Figure S11. Full LLM interpretation of the decision tree for Ki-67 expression control.** The model identifies cellular conditions with high PTMA attribution and proposes biological rationales, testable hypotheses, and supporting literature based on the decision tree logic. It suggests that a sequence of post-transcriptional regulatory events, coupled with PTMA-associated chromatin remodeling, could drive Ki-67 expression.

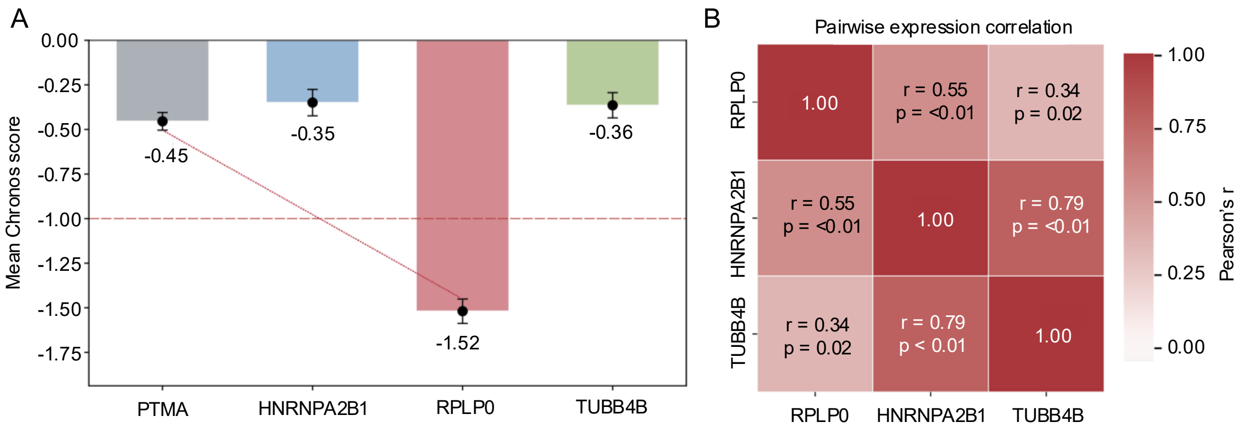


**Figure S12. Orthogonal validation of the LLM-hypothesized Ki-67 hierarchical regulatory axis.** (A) Evaluation of genetic essentiality utilizing mean Chronos scores. The downstream effector *RPLP0* demonstrates significant essentiality, mapping well below the threshold (Chronos < -1). Conversely, the upstream initiator *PTMA*, along with *HNRNPA2B1* and the mitotic marker *TUBB4B*, exhibit milder essentiality (Chronos = -0.45, -0.35 and -0.36, respectively). (B) Pairwise expression correlations examining the relationship between post-translational processors and mitotic activity. *HNRNPA2B1* expression demonstrates statistically significant positive correlations with both the mitotic marker *TUBB4B* (Pearson's r = 0.79, p < 0.01) and the translation effector *RPLP0* (r = 0.55, p < 0.01), supporting functional coupling and corroborating the discrete subpopulation logic. *n* = 51 breast cancer genome-wide CRISPR knockout models.

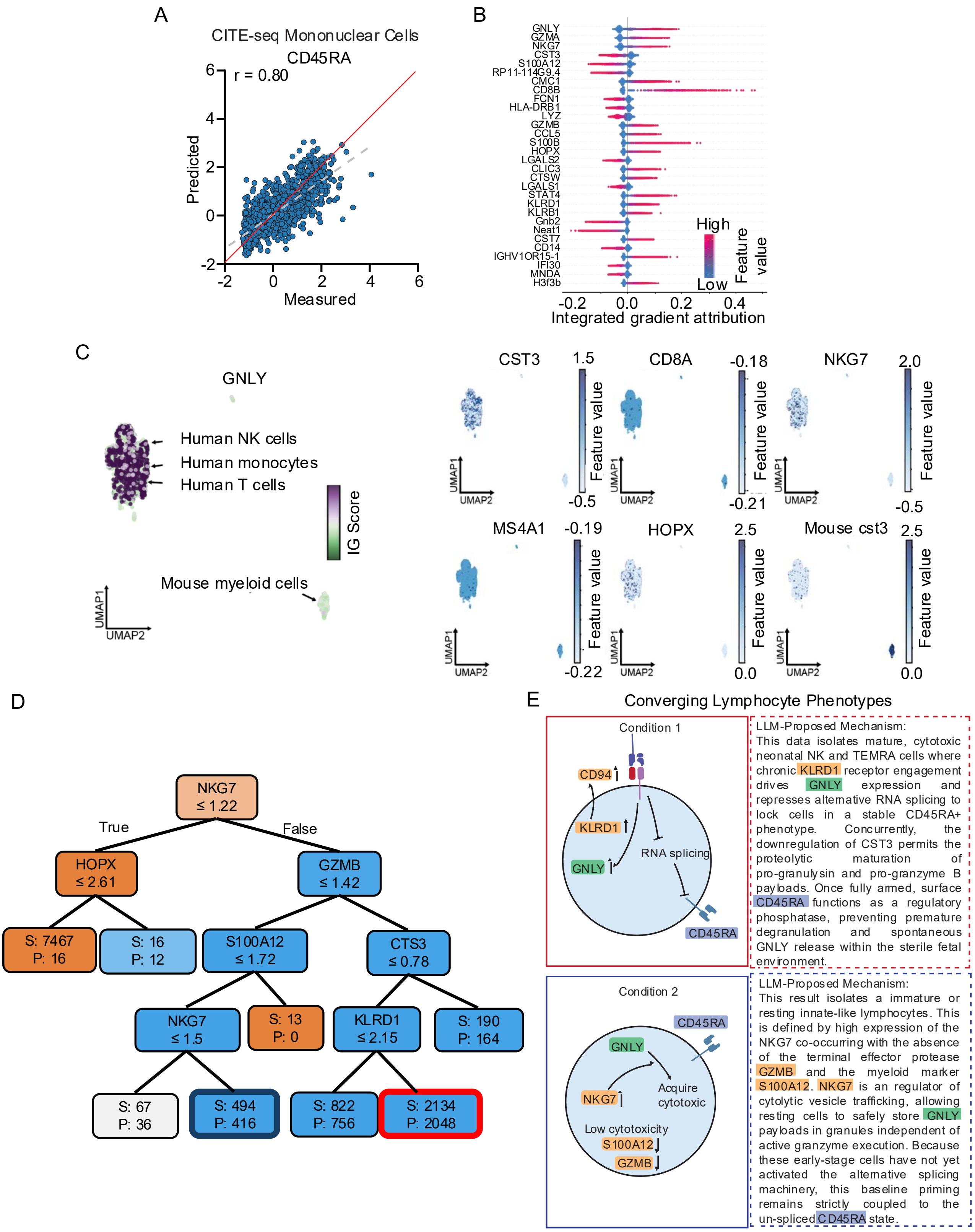


**Figure S13. PITCH predicts convergent CD45RA+ phenotypes.** (A) The MLP accurately predicts naïve immune marker CD45RA using single-cell RNA profile (Pearson's r= 0.80, gray dash line; low residual error, red identity line). (B) IG score ranking shows that the associated features are primarily cytotoxic effector genes (e.g., *GZMA*, *NKG7*, *GZMB*), identifying *GNLY* as the primary predictor. (C) UMAP projection of *GNLY* IG scores and RNA markers shows that canonical marker expression but fails to fully resolve immune lineages, limiting direct interpretation of *GNLY* IG score distribution. (D) The DT isolates a convergent *GNLY*$^{+}$/CD45RA$^{+}$ phenotype characterized by high-*NKG7*, high-*GZMB*, low-*CST3*, and high-*KLRD1* expression (Condition 1, red frame). A second subpopulation is defined by high-*NKG7*, low-*GZMB*, low-*S100A12* (Condition 2, blue frame). (E) LLM's hypotheses for the two subpopulations associated with *GNLY*-linked CD45RA expression: a mature NK-like cytotoxic population ((D), Condition 1) and a primed cytotoxic lymphocyte population ((D), Condition 2).

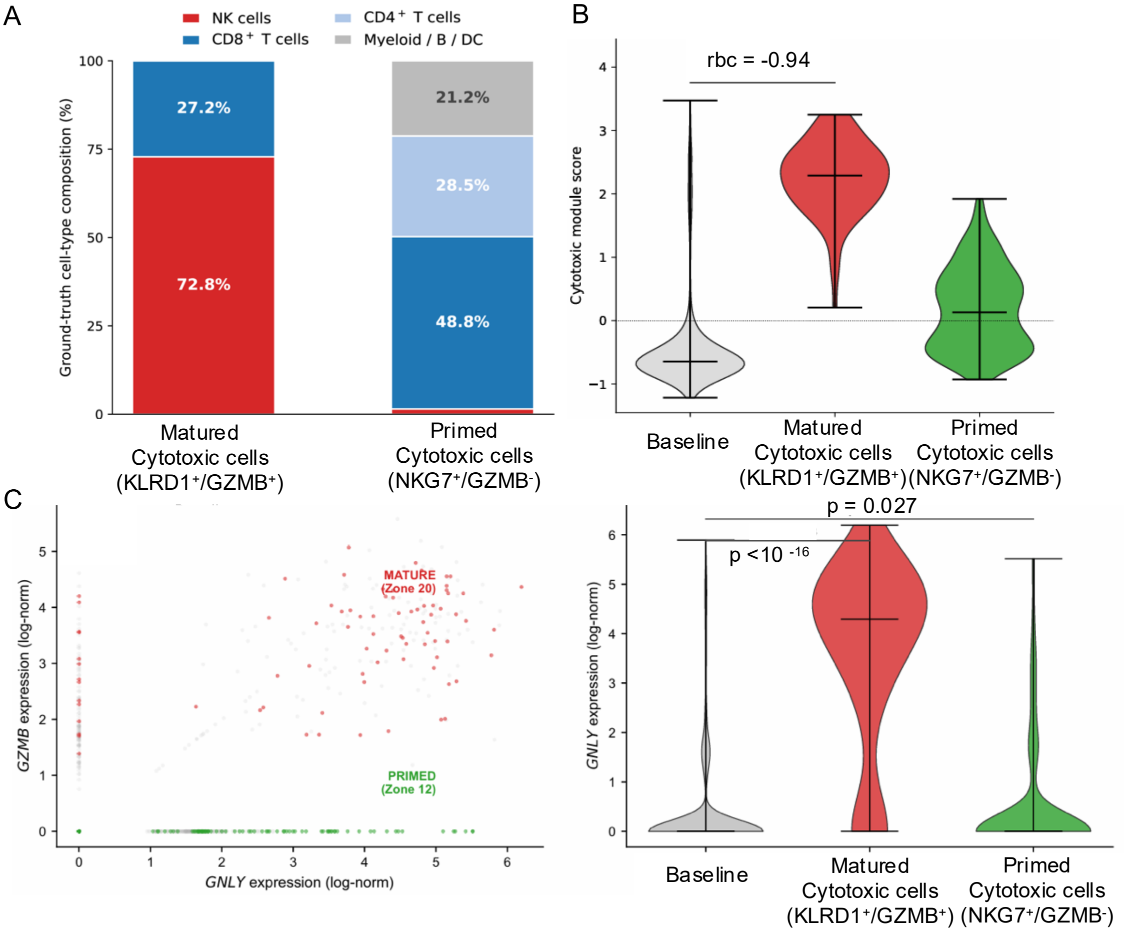


**Figure S14. Validation of convergent CD45RA-associated cytotoxic lymphocyte phenotypes.** (A) Cell-type composition of the PITCH-defined gates in an independent PBMC3K cohort. The *KLRD1*$^+$/*GZMB*$^+$ gate is enriched for cytotoxic lineages (NK and CD8$^+$ T cells), whereas the *NKG7*$^+$/*GZMB*$^-$ gate is dominated by CD8$^+$ T cells with additional CD4$^+$ and remain blood cells. (B) Cytotoxic module scores validate the predicted matured versus primed cell lineages : *KLRD1*$^+$/*GZMB*$^+$ cells show strong cytotoxic elevation (rbc = -0.94), while *NKG7*$^+$/*GZMB*$^-$ cells show a weaker but detectable cytotoxic state. (C) Validation of LLM-proposed *GNLY*-*GZMB* decoupling hypothesis. In primed cytotoxic cells, the expression of *GNLY* is independent to terminal cytotoxic effector *GZMB*, while *GNYL* and *GZMB* are co-expressed in mature cytotoxic cells (left). The violin plot of *GNLY* shows that *GNYL* is elevated in both conditions compared to the baseline ($p < 10^{-16}$ for matured cytotoxic cells and $p = 0.027$ for primed cytotoxic cells, right), suggesting that *GNLY* upregulation can precede terminal *GZMB*-associated cytotoxic maturation.

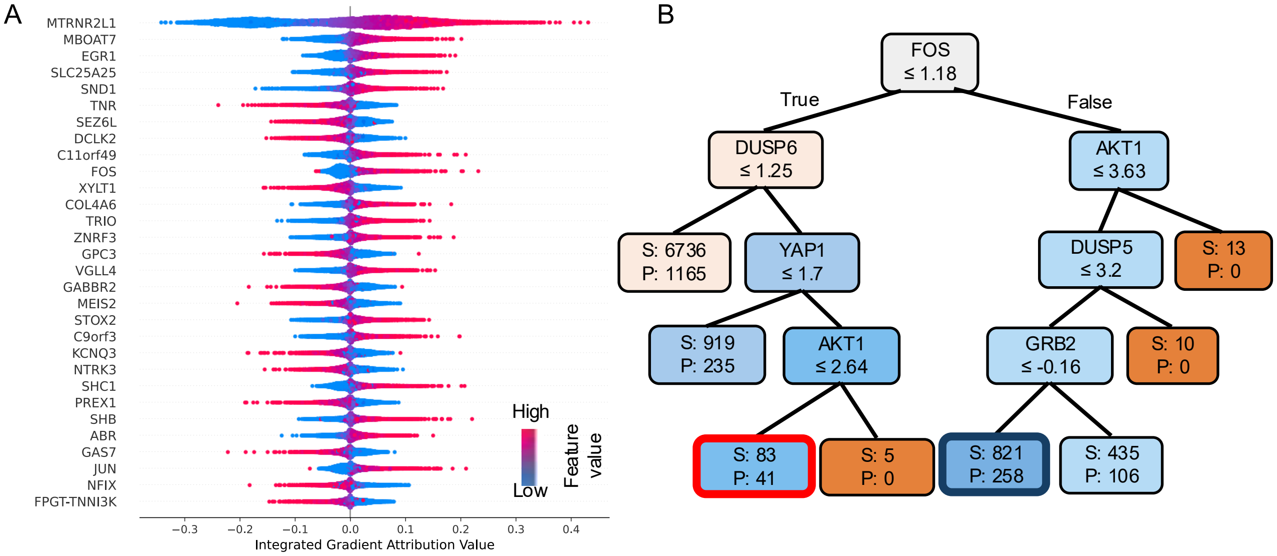


C

Condition 1:

Biological Rationale:
This zone describes a cellular state where canonical PI3K/AKT signaling is constrained (AKT1 is low) and upstream receptor tyrosine kinase (RTK) signaling is dampened (GRB2 is low). However, immediate early gene (IEG) transcription is highly active (FOS is high). Crucially, DUSP5, a nuclear phosphatase that specifically dephosphorylates and deactivates ERK, is low. Because AKT1 is low, the mTORC1-S6K axis is likely inactive. Therefore, pRPS6 must be driven by the alternative MAPK/ERK -> p90RSK -> pRPS6 pathway. The low levels of DUSP5 allow for sustained nuclear ERK activity. Sustained ERK is known to drive the robust transcription of EGR1. In this state, EGR1 acts as the ultimate surrogate marker for a hyperactive, sustained ERK/RSK signaling axis that directly phosphorylates RPS6.

Confirmations of Literature Knowledge:
ERK/RSK-mediated RPS6 Phosphorylation: It is well established that pRPS6 can be phosphorylated independently of AKT/mTOR via the ERK-p90RSK pathway.DUSP5 as a Nuclear ERK Regulator: DUSP5 specifically anchors to and deactivates ERK in the nucleus. Low DUSP5 correlates with prolonged nuclear ERK signaling, which is required for the transcription of IEGs like FOS and EGR1.2.

Hypothesis:
In developing neural organoids, transient RTK signaling (low GRB2) triggers a sustained nuclear ERK response (due to low DUSP5 feedback) that drives protein synthesis (pRPS6) independently of AKT. EGR1 is not just a bystander but may actively transcribe secondary targets required to sustain this AKT-independent ribosomal activity.

Condition 2:

Biological Rationale:
This zone describes a cellular state where canonical MAPK/ERK signaling is actively suppressed: FOS is low, and DUSP6 (a cytoplasmic ERK-specific phosphatase) is high. AKT1 remains low. Surprisingly, YAP1 (the primary effector of the Hippo pathway) is highly expressed. How does EGR1 predict pRPS6 when both AKT and ERK pathways appear suppressed? YAP1 is a mechanosensitive transcription factor critical for neural progenitor expansion. Literature suggests that YAP1 can bypass canonical RTK signaling to activate mTORC1 (via amino acid transporters or PTEN suppression) and can directly co-regulate genes with TEAD. In this state, EGR1 expression is likely driven by YAP1/TEAD mechanotransduction rather than ERK. EGR1 here serves as a biomarker for YAP1-mediated cellular growth and translation (pRPS6) in neural progenitors experiencing specific physical/mechanical cues within the 3D organoid environment.

Confirmations of Literature Knowledge:
YAP1 in Neural Organoids: YAP1 is highly active in the ventricular zone of brain organoids, driving the proliferation of radial glia and neural progenitors.DUSP6 as an ERK Feedback Loop: High DUSP6 is a classic negative feedback mechanism that shuts down cytoplasmic ERK, correlating perfectly with the low FOS seen in this node.

Hypothesis: In the 3D architecture of brain organoids, mechanical stress activates YAP1, which compensates for the lack of canonical growth factor signaling (low AKT, low ERK/high DUSP6) to maintain protein synthesis (pRPS6). YAP1 directly drives EGR1 transcription, and together they maintain the translation machinery required for neural progenitor expansion.

**Figure S15. Retrospective reconstruction of AKT-independent pRPS6 regulatory axes by PITCH.** (A) IG rank identifies importance transcriptomic features associated with pRPS6 abundance, with top positive predictors dominated by immediate-early transcription factors (e.g., *EGR1*, *FOS*, *JUN*). (B) Applying pRPS6 regulatory features to classify high *EGR1* IG score, the DT uses early MAPK effector FOS as the root split to resolve two regulatory axes: a YAP1/TEAD-associated subpopulation (Condition 1, red frame) and a MAPK-dependent pRPS6 regulatory subpopulation (Condition 2, blue frames). (C) Full rationales and hypotheses proposed by the LLM based on the two subpopulations logic in (B). LLM hypothesizes that pRPS6 regulation can be PI3K/AKT-independent and driven by MAPK/ERK signaling where *FOS* is high (Condition 1, red frame), or associated with YAP1/TEAD-mediated program when *FOS* is low (Condition 2, blue frame).

A

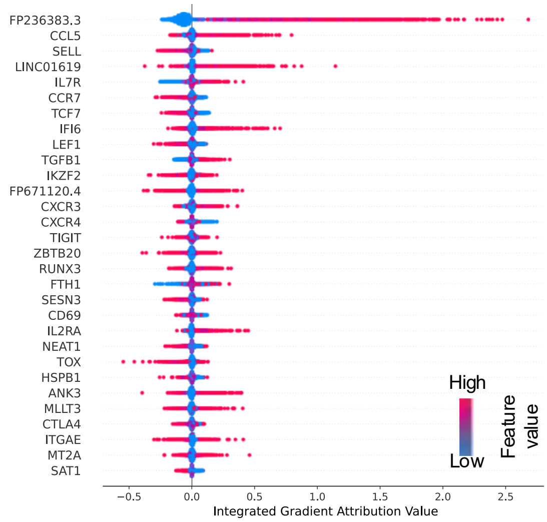


B

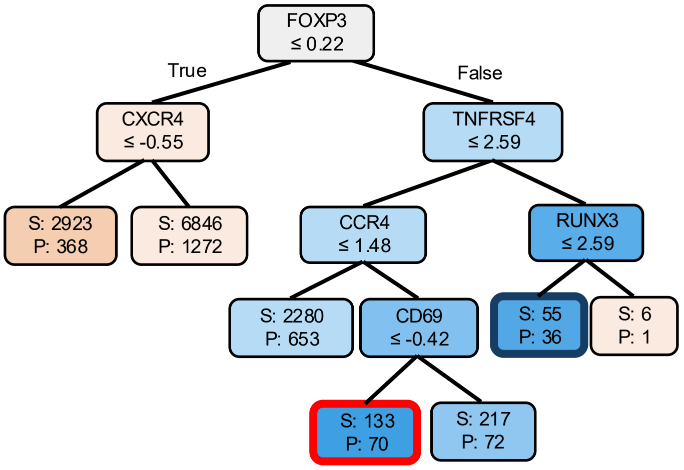


C

Condition 1:

Biological Rationale:
This node captures a highly specific population of Tregs (FOXP3+) that are actively infiltrating the tumor but have not yet established tissue residency. CCR4 is the classic chemokine receptor used by Tregs to migrate into the tumor microenvironment (TME) along CCL22 gradients produced by tumor-associated macrophages. The lack of CD69 (a classic marker of tissue-resident memory T cells, Trm, and early activation) and moderate/low TNFRSF4 (OX40, a marker of terminal Treg activation) suggests these are newly arrived, circulating-to-infiltrating Tregs. In this specific transitional state, IL2RA is highly predictive of CCR5. This implies that as these CCR4+ Tregs enter the TME, they rely heavily on IL-2 signaling (via IL2RA) to upregulate CCR5, which is necessary to navigate deeper into the inflammatory, CCL5-rich core of the HCC tumor.

Findings Confirming Literature Knowledge:
FOXP3 and CCR4 co-expression: It is well-established that CCR4 is a primary driver of Treg recruitment into solid tumors, including HCC. IL-2 driving chemokine receptor expression: IL-2 signaling is known to be required for the differentiation of resting Tregs into effector Tregs (eTregs), a process that includes the upregulation of CCR5 to allow migration to sites of inflammation.

Hypothesis:
In HCC, newly recruited Tregs (CCR4+, CD69-) require an initial pulse of IL-2 (captured by high IL2RA attribution) to undergo a spatial "handoff" from CCR4-mediated entry to CCR5-mediated deep-tumor retention.

Condition 2:

Biological Rationale:
While Node 21 represents newly arriving Tregs, Node 25 represents a highly activated, terminally differentiated effector Treg population. TNFRSF4 encodes OX40, a costimulatory molecule that, when highly expressed on Tregs in the TME, marks a hyper-suppressive state associated with poor prognosis in HCC. RUNX3 is a transcription factor that typically promotes cytotoxic (Th1/CD8) lineages; its downregulation is actively required to maintain FOXP3 stability and prevent Tregs from converting into "fragile" ex-Tregs that produce IFN-gamma. In this highly stable (RUNX3 low), highly activated (TNFRSF4 high) state, IL2RA perfectly predicts CCR5. This suggests that in the most suppressive Tregs, OX40 costimulation synergizes with IL-2 signaling to lock in the CCR5+ phenotype, ensuring these cells remain anchored in the tumor bed.

Findings Confirming Literature Knowledge:
OX40 (TNFRSF4) as an eTreg marker: High OX40 expression is a hallmark of highly suppressive, tumor-infiltrating Tregs in HCC. RUNX3 and FOXP3 antagonism: Literature confirms that RUNX3 binds to the Foxp3 locus to repress it. Therefore, low RUNX3 is a biological prerequisite for maintaining high FOXP3 expression and Treg lineage stability in inflammatory environments.

Hypothesis:
OX40 signaling (TNFRSF4 > 2.590) and RUNX3 suppression (RUNX3 <= 2.058) create a synergistic transcriptional environment that makes CCR5 expression entirely dependent on continuous IL2RA (IL-2) signaling in HCC-infiltrating Tregs.

**Figure S16. Retrospective reconstruction of Treg cell maturation by PITCH.** (A) A beeswarm plot of IG scores illustrates that the top-ranked transcripts are associated with immune cell localization (e.g., *CCL5*, *CCR7*) and T cell regulatory states (e.g., *IL2RA*, *IKZF2*). (B) Applying Treg status features to classify high *IL2RA* IG score, the tree identified both Treg subsets are *FOXP3*$^+$. A subsequent split based on the maturation marker *TNFRSF4* delineates a less mature, high-*CCR4* transitional state (Condition 1, red frame) from a highly mature, *TNFRSF4*-enriched subset (Condition 2, blue frame). (C) Full rationales and hypotheses generated by the LLM for the two *FOXP3*$^+$ Treg states identified in (B). The model distinguishes between a transitional state actively infiltrating the tumor via a *CCR4*-to-*CCR5* handoff (Condition 1, red frame) and a terminally differentiated, tumor-resident effector state reliant on *IL-2* signaling (Condition 2, blue frame).

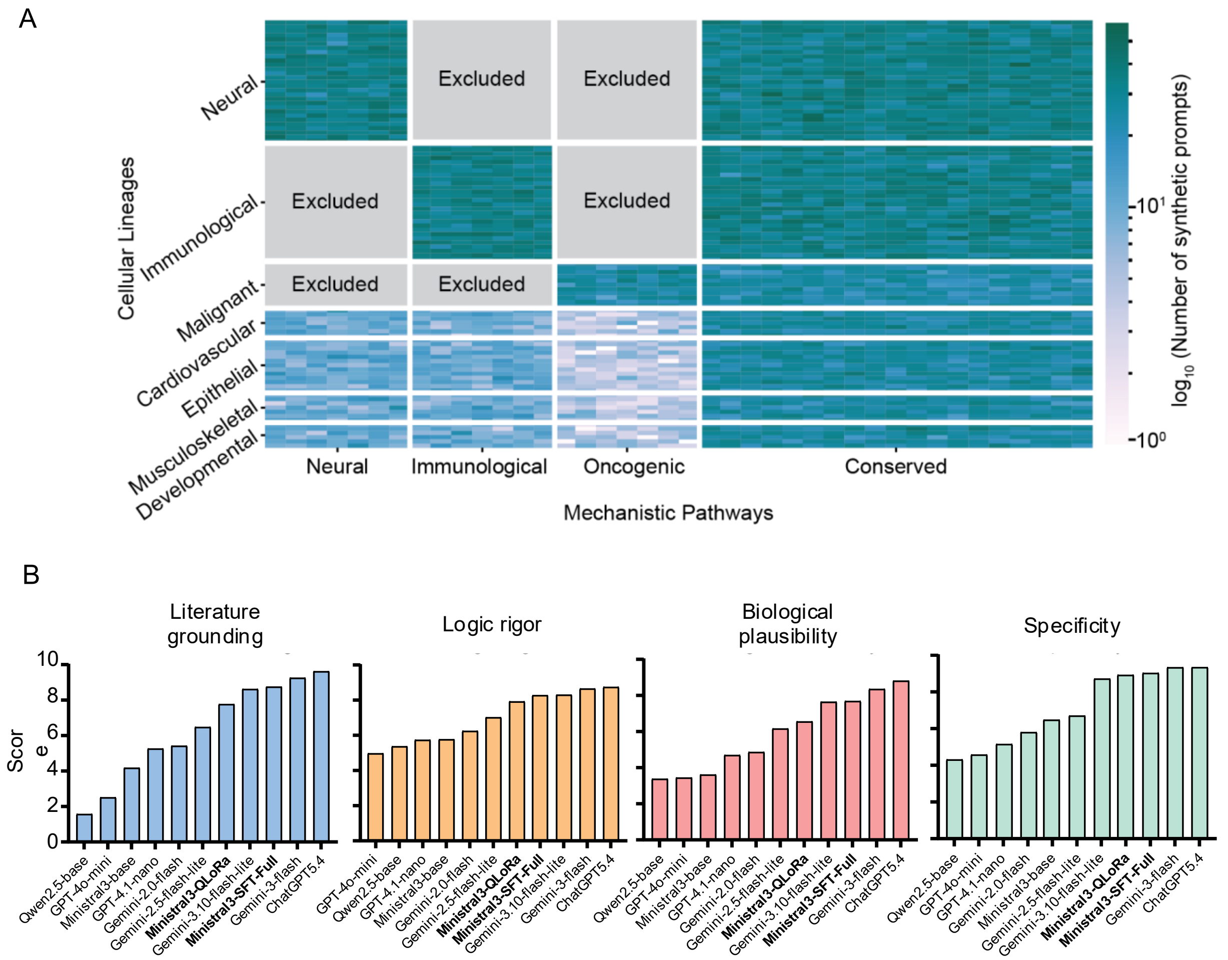


**Figure S17. Training and evaluation of local LLMs for DT interpretation.** (A) A total of 30,000 decision tree prompts were synthesized and randomized to encompass broad biological contexts. To ensure LLM training was conducted on self-consistent data, synthetic trees derived from biologically incompatible combinations of contexts (gray boxes) were excluded. (B) Benchmarking of the local Ministral 3-SFT model against industry-standard LLMs across four evaluation pillars shows that fine-tuning substantially improves performance over the base architecture. The fine-tuned model surpasses the economy-tier Gemini 3.10 Flash-Lite model and achieves performance comparable to Gemini 1.5 Flash in tree interpretation.

| Feature name |
|---|
| Nucleus Morphology Area |
| Nucleus Morphology Perimeter |
| Nucleus Morphology Elongation |
| Morphology Area |
| Morphology Perimeter |
| Morphology Elongation |
| Cytoplasm Morphology Area |
| Cytoplasm Morphology Perimeter |
| Cytoplasm Morphology Elongation |
| Membrane Morphology Area |
| Membrane Morphology Perimeter |
| Membrane Morphology Elongation |
| Cytobody Morphology Area |
| Cytobody Morphology Perimeter |
| Cytobody Morphology Elongation |
| Nucleus Intensity mean DAPI |
| Cytoplasm Intensity mean Paxillin |
| Nucleus Intensity mean YAP |
| Membrane Intensity mean YAP |
| Nucleus Intensity mean PCNA |
| Cytobody Intensity mean Paxillin |
| Nucleus Intensity std DAPI |

**Table S1. List of mechanobiology features.** The 22 mechanobiology features curated for training decision tree target cells with high Actin attribution to pERK activity.

| Feature name | | | | |
|---|---|---|---|---|
| **Immediate-early response markers** | **RTK / upstream adaptor module** | **ERK / MAPK signaling branch** | **AKT-mTOR-S6K branch** | **YAP/TEAD regulatory program** |
| FOS | SHC1 | RPS6KA1 | RPS6KB1 | TEAD1 |
| JUN | SHB | MAPK1 | MTOR | TEAD3 |
| JUNB | GRB2 | MAPK3 | AKT1 | YAP1 |
| | SOS1 | MAP2K1 | | VGLL4 |
| | EGFR | MAP2K2 | | |
| | | DUSP1 | | |
| | | DUSP4 | | |
| | | DUSP5 | | |
| | | DUSP6 | | |

**Table S2. List of pRPS6 regulatory features.** The RNA features for immediate-early transcriptional response, RTK/upstream adaptor signaling, ERK/MAPK signaling, AKT-mTOR-S6K signaling and YAP/TEAD regulatory program are selected to investigate the RPS6 phosphorylation in neuron.

| Feature name | | | |
|---|---|---|---|
| **Treg Identity & Suppression** | **Chemokine-Mediated Recruitment** | **Tumor-Resident Maturation** | **Tumor-Adapted Activation** |
| FOXP3 | CCR8 | LAYN | TOX |
| CTLA4 | CCR5 | CD69 | BATF |
| TIGIT | CCL5 | ITGAE | IRF4 |
| ICOS | CCR4 | PRDM1 | PDCD1 |
| TNFRSF18 | CCR6 | RUNX3 | |
| TNFRSF4 | CXCR3 | ZNF683 | |
| IKZF2 | CXCR4 | | |
| | CXCR6 | | |

**Table S3. List of Treg status features.** The RNA features of Treg identity and suppressive, migration and chemokine-axis markers, tumor-resident maturation markers and tumor adaptation markers are used in training decision tree targeting cells with high IL2RA-CCR5 protein association.

| Axis | Configuration | # runs | Pearson r | $R^2$ | RMSE |
|---|---|---|---|---|---|
| **Default MLP** | Default: h = 256, dropout= 0.5, wd = $2 \times 10^{-3}$ † | 5 folds | $0.904 \pm 0.006$ | $0.814 \pm 0.011$ | $0.674 \pm 0.022$ |
| **Architecture** | hidden = 32 | 5 folds | $0.846 \pm 0.008$ | $0.627 \pm 0.016$ | $0.955 \pm 0.018$ |
| | hidden = 128 | 5 folds | $0.894 \pm 0.006$ | $0.792 \pm 0.010$ | $0.712 \pm 0.018$ |
| | hidden = 256 † | 5 folds | $0.904 \pm 0.006$ | $0.814 \pm 0.011$ | $0.674 \pm 0.022$ |
| | hidden = 512 | 5 folds | $0.906 \pm 0.006$ | $0.819 \pm 0.011$ | $0.664 \pm 0.023$ |
| | hidden = 1024 | 5 folds | $0.905 \pm 0.007$ | $0.818 \pm 0.013$ | $0.666 \pm 0.025$ |
| **Regularization** | dropout= 0, weight decay= 0 | 5 folds | $0.874 \pm 0.004$ | $0.757 \pm 0.008$ | $0.771 \pm 0.016$ |
| | dropout= 0.25, weight decay= 0 | 5 folds | $0.893 \pm 0.007$ | $0.794 \pm 0.014$ | $0.710 \pm 0.025$ |
| | dropout= 0.5, weight decay= 0 | 5 folds | $0.895 \pm 0.007$ | $0.797 \pm 0.014$ | $0.703 \pm 0.026$ |
| | dropout= 0, weight decay= $2 \times 10^{-3}$ | 5 folds | $0.888 \pm 0.004$ | $0.784 \pm 0.008$ | $0.728 \pm 0.017$ |
| | dropout= 0.25, weight decay= $2 \times 10^{-3}$ | 5 folds | $0.900 \pm 0.006$ | $0.808 \pm 0.012$ | $0.685 \pm 0.023$ |
| | dropout= 0.5, weight decay= $2 \times 10^{-3}$ † | 5 folds | $0.904 \pm 0.006$ | $0.814 \pm 0.011$ | $0.674 \pm 0.022$ |
| **Random seed** | MLP best config (h = 512, dropout= 0.5, weightdecay = $2 \times 10^{-3}$) | 10 seeds | $0.908 \pm 0.003$ | $0.823 \pm 0.006$ | $0.658 \pm 0.014$ |

**Table S4. Robustness and hyperparameter sensitivity of the 2-layer MLP.** The predictive performance of the MLP for pERK prediction at 10 ng/mL EGF was evaluated across varying architectural and regularization configurations using five-fold cross-validation. The symbol † indicates the default optimized configuration. Performance improvements associated with the increasing the hidden layer nodes reach a plateau at 256. Additionally, the inclusion of the dropout and weight decay is shown to be critical for model regularization. All values represent mean ± standard deviation.

| Rank | Model | Pearson r | RMSE |
|---|---|---|---|
| 1 | MLP | $0.904 \pm 0.006$ | $0.674 \pm 0.022$ |
| 2 | LightGBM | $0.892 \pm 0.004$ | $0.709 \pm 0.015$ |
| 3 | HistGradientBoosting | $0.890 \pm 0.003$ | $0.716 \pm 0.011$ |
| 4 | SVR (RBF) | $0.869 \pm 0.006$ | $0.807 \pm 0.018$ |
| 5 | Random Forest | $0.826 \pm 0.006$ | $0.919 \pm 0.012$ |
| 6 | KNN (k = 20) | $0.664 \pm 0.008$ | $1.217 \pm 0.010$ |

**Table S5. Five-fold cross-validation benchmark of the MLP against representative nonlinear models.** Six non-linear regressors were evaluated using 10 ng/mL EGF stimulation data, comprising $n$ = 23,569 samples with 510 features. Pearson r and RMSE are reported as mean ± standard deviation across the five outer folds. The MLP architecture attains the highest Pearson r = 0.904 and lowest RMSE = 0.674.

| Model | Pathway Hallucinatio (%) | Context Mismatch (%) | Causal Overreach (%) | Logic Failure (%) | Vagueness (%) |
|---|---|---|---|---|---|
| Ministral 3-QLoRa | 31.2 | 33.2 | 21.6 | 1.0 | **0.0** |
| Ministral 3-SFT | 23.6 | 21.6 | 19.1 | 1.0 | **0.0** |
| Ministral 3-Base | 70.9 | 55.3 | 64.3 | 5.5 | **0.0** |
| Gemini 3.0 Flash | 12.1 | 15.6 | 9.5 | 0.5 | **0.0** |
| ChatGPT 5.4 | **0.5** | **14.1** | **1.0** | **0.0** | **0.0** |
| Gemini 3.1 Flash-Lite | 16.6 | 21.1 | 16.6 | 0.5 | **0.0** |
| Gemini 2.5 Flash-Lite | 18.6 | 32.2 | 19.1 | 2.5 | 11.1 |
| Gemini 2.0 Flash | 28.1 | 49.2 | 42.2 | 1.5 | 19.1 |
| ChatGPT 4o-mini | 57.8 | 58.8 | 61.8 | 8.0 | 48.7 |
| ChatGPT 4.1-nano | 37.2 | 56.3 | 53.3 | 5.0 | 33.7 |
| Qwen 2.5-Base | 49.7 | 64.8 | 55.3 | 2.0 | 65.3 |

**Table S6. Benchmarking the failure rates of LLMs in interpreting synthetic tree prompts.** Knowledge distillation from the Gemini 3.1 Pro significantly reduces the failure rates of student model interpretation. Generate detailed interpretation (Vagueness = 0 %) and plausible hypothesis comparable to Gemini 3.1 Flash-Lite. ChatGPT 5.4 as a positive control confirms the robustness of the criteria by exhibiting the lowest failure rates across all five metrics.
